# Origins of reactivity in SAM-utilizing ribozyme SAMURI-catalyzed RNA alkylation

**DOI:** 10.64898/2026.04.24.720726

**Authors:** Julie Puyo-Fourtine, Yanan Du, Erika McCarthy, Şölen Ekesan, Darrin M. York

## Abstract

Unlocking the design principles of programmable RNA catalysts capable of sitespecific chemical modification is critical for expanding the functional and therapeutic potential of RNA. The SAM analogue-utilizing ribozyme (SAMURI) enables sitespecific RNA alkylation using either S-adenosylmethionine (SAM) or the synthetic cofactor propargylic Se-2,6-diaminopurinribosyl-selenomethionineamide (ProSeDMA), yet the molecular determinants of its reactivity remain incompletely understood. Here, we combined molecular dynamics, 3D-RISM solvation analysis, alchemical free-energy calculations, quantum p*K_a_* shift predictions, and *ab initio* QM/MM free-energy simulations to characterize the conformational and electronic factors that govern catalysis. Simulations show that, although the global fold of SAMURI remains stable in solution, the formation of catalytically competent near-attack configurations is rare, indicating that the observed rate depends on access to a minor fraction of these reactive conformations (*f_react_*). A putative Mg^2+^ binding site between the SAM carboxylate and the G30 phosphate, together with a hydrogen bond between the cofactor *α*-amine and U8:O2, enriches *f_react_*. QM/MM simulations support an S_N_2-like alkyl transfer mechanism and show that ProSeDMA reacts more readily than SAM primarily due to its more favorable electronic leaving-group properties that enhance the intrinsic rate (*k_int_*). Atomic substitutions at A52 that tune the N3 p*K_a_* enhance nucleophilicity, further lower the activation barrier, and increase *k_int_*. Together, these results show that SAMURI catalysis is governed by a combination of conformational preorganization and electronic effects, providing a framework to guide the design of new programmable RNA alkyltransferases.

**TOC Graphic:** 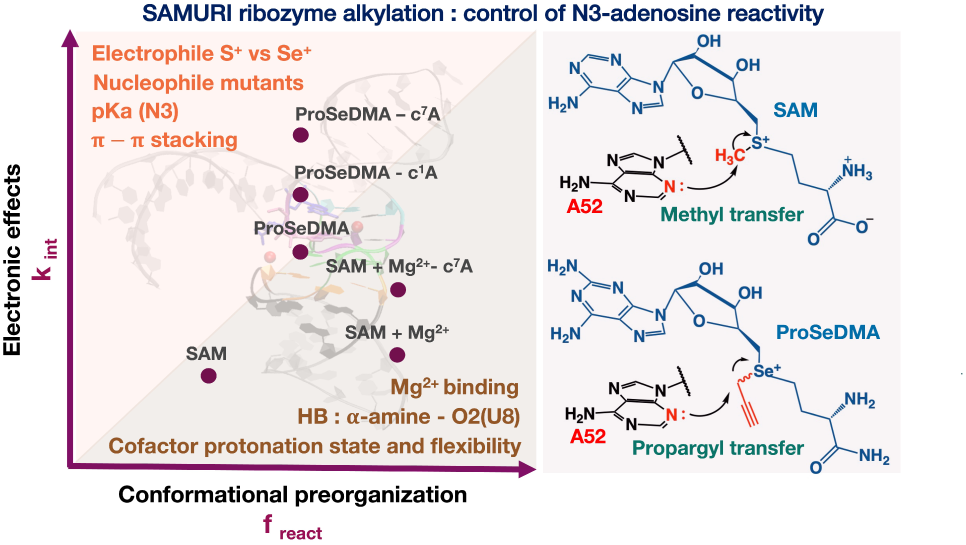

## Introduction

Nucleic acids can fold into complex three-dimensional architectures with catalytic activity, enabling them to function as enzymes in the absence of proteins, the so-called *ribozymes*.^1^ Their discovery triggered a paradigm shift by overturning the long-standing view that only proteins could act as biological catalysts^2–4^ and provided strong support for the hypothesis that RNA was the primary self-replicating biomolecule in early life. ^5,6^ Across all kingdoms of life, naturally occurring ribozymes participate in essential biological processes such as protein synthesis in the ribosome and tRNA maturation.^7,8^ Many of them exhibit sequence-specific RNA cleavage activity that can be exploited for therapeutic applications, for instance in anticancer strategies targeting tumor-specific RNAs, as well as in biosensing platforms.^9,10^

Beyond their biological roles, ribozymes provide key evidence supporting the RNA world hypothesis.^5,6^ In this scenario, early self-replicating systems were composed entirely of RNA, preceding the emergence of protein synthesis and DNA-based information storage. However, natural ribozymes display a limited catalytic repertoire, acting predominantly as nucleolytic catalysts that promote self-cleavage or ligation reactions, and are largely restricted to phosphoryl transfer, with only a few examples extending to peptide bond formation.^11–14^ A broader range of reactions would nevertheless have been required to sustain a primitive metabolism, including C–C and C–N bond formation essential for the synthesis of molecular precursors, yet are absent in known natural ribozymes.^6,15–17^ Artificial ribozymes with such activities therefore provide a powerful strategy to explore plausible scenarios for early metabolic evolution by probing the chemical potential of RNA.

In the early 1990s, it was demonstrated that catalytic RNAs could be evolved toward new activities through iterative cycles of mutation, selection and amplification. This *in vitro* approach, inspired by aptamer discovery, starts from large libraries of randomized RNA sequences challenged with a specific chemical task and enriches rare active sequences through successive rounds of selection.^18^ Over the past three decades, these methods have yielded ribozymes capable of RNA ligation and polymerization,^19,20^ as well as catalyzing small-molecule transformations, including Diels–Alder cycloadditions,^21^ aldol condensations,^22^ Michael additions^23^ and site-specific alkylations.^9,24–27^ These findings demonstrate that RNA catalysis extends far beyond what has been so far observed in nature.

Among these reactions, alkyl group transfer is of particular biological and technological interest. In living systems, alkylation of nucleic acids is a pervasive modification that can modulate base-pairing, alter charge and hydrogen-bonding patterns, influence RNA folding and regulate interactions with proteins. ^28^ In tRNA and rRNA, specific methylations help stabilize structure and support accurate translation. ^29,30^ More generally, alkyl transfer reactions enable the introduction of reactive handles for fluorescence labeling, affinity capture or chemical modification.^31–33^

Such reactions are predominantly catalyzed by protein methyltransferases using S-adenosylmethionine (SAM) as the alkyl donor.^34^ However, these enzymes typically recognize conserved sequence motifs or structural folds, making site-specific RNA modification challenging.^25,35^ Several studies have demonstrated that RNA can be evolved to bind an alkyl donor and promote selective transfer, including the self-biotinylating ribozyme,^9^ the SAM-dependent guanine N7 methyltransferase SMRZ-1,^25^ the *O*^6^-alkylguanine–dependent adenine N1 alkyltransferases MTR1 and RACR^24,26,36^ and the *O*^6^-benzylguanine-depedent cytosine N4 alkyl transferase CSAR.^26,27^

Recently, the SAM analogue-utilizing ribozyme (SAMURI) was shown to form a fully RNA-based active site capable of site-specific alkylation.^31,37^ Evolved *in vitro* from an RNA library, this ribozyme uses a chemically stabilized and bioorthogonal analogue of SAM; propargylic Se-2,6-diaminopurinribosyl-selenomethionineamide (ProSeDMA), to transfer a propargyl group to the N3 position of a specific adenosine in a target RNA. The resulting alkyne provides a small reactive handle for click chemistry, allowing subsequent labeling with fluorophores, affinity tags and related probes.^33^ SAMURI remains active at physiological Mg^2+^ concentrations and retains catalytic activity in mammalian cells, making direct intracellular RNA labeling possible. It can also use natural SAM to produce N3-methyladenosine (m^3^A), extending its reactivity beyond the engineered cofactor.

Structural studies^31,37^ showed that SAMURI adopts a compact three-helix junction in which the catalytic core is organized into four stacked layers that preorganize the reactive groups for chemistry. The upper layer contains the donor ProSeDMA in a base triple, where stacking interactions and hydrogen bonds orient the reactive selenium center for nucleophilic attack. In the next layer, the target adenosine A52 is positioned in a second base triple between P1 and P3, aligning its N3 atom with the cofactor in an S_N_2 -like arrangement. Below this, two base pairs stabilize the fold, and a lower layer further reinforces the stack. Two hydrated magnesium ions also contribute to the organization of the junction by linking backbone phosphates and supporting the kink that inserts A52 into the active site. More generally, prior structural and computational studies have shown that metal ions and their hydration shells play central roles in organizing RNA folds and shaping catalytically relevant conformations.^38–40^ This architecture helps explain both the efficiency and the selectivity of SAMURI by juxtaposing the alkyl donor with the reactive nucleophile while disfavoring unproductive geometries (Figure 1).

**Figure 1:**
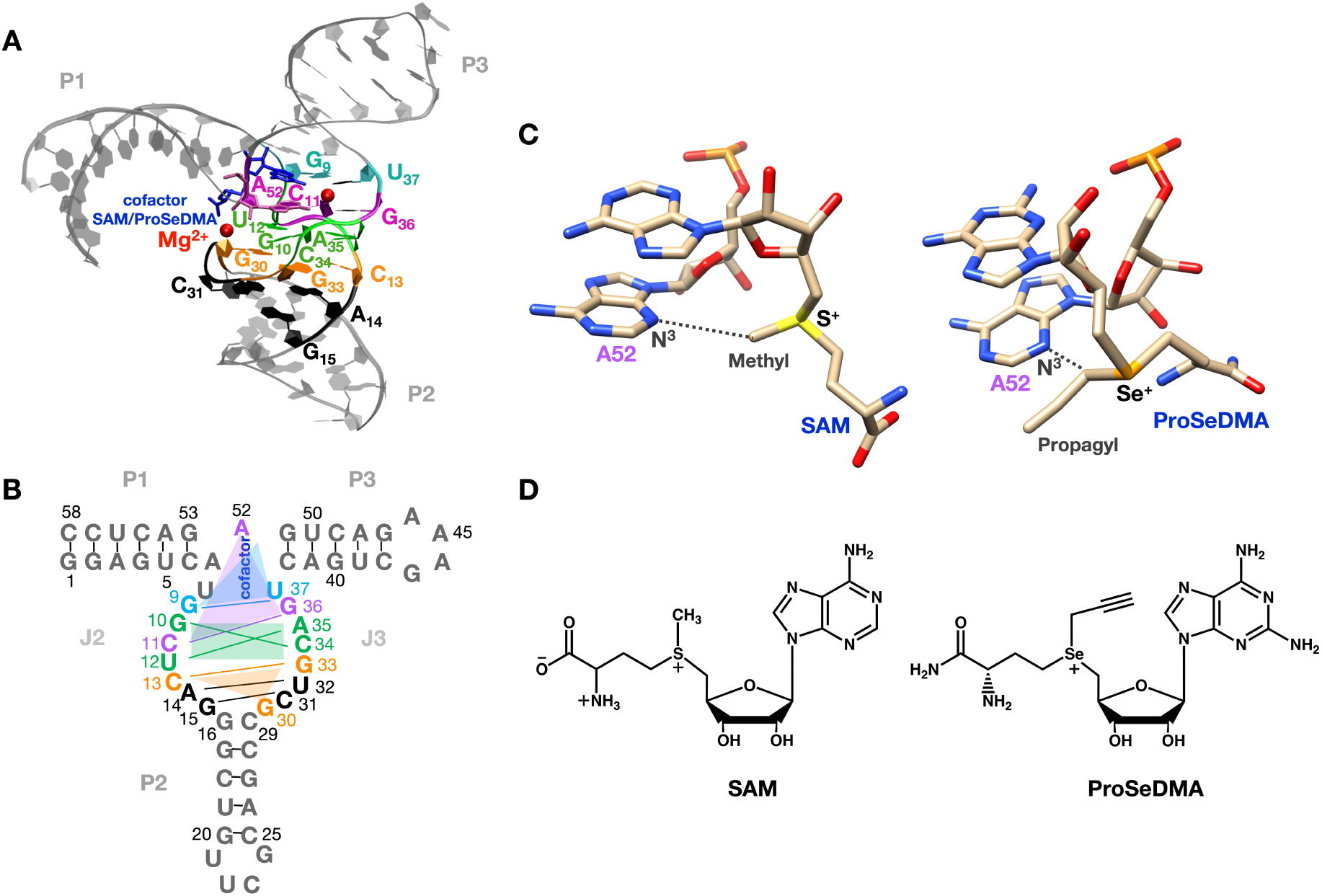
Structure of the SAMURI ribozyme. **(A)** Representation of SAMURI with the catalytic core highlighted by its four stacked layers: the *cofactor layer* (blue) defined by a base triple, the *reaction layer* (magenta) where the target adenosine (A52) engages in a base triple, the *stabilizing layer* (green) composed of two coplanar Watson–Crick base pairs and the *bottom layer* (orange) formed by another base triple. The zipper region is shown in black, while helices P1, P2, P3 and loop regions are in grey. **(B)** Secondary structure of the 58-nt SAMURI construct, colored according to the scheme in **(A)**. **(C)** Active site views with the two cofactors used in this study: S-adenosylmethionine (SAM, left) enabling methyl transfer, and the synthetic analog Propargylic Se-2,6-diaminopurin-ribosylselenomethionineamide (ProSeDMA, right) enabling propargyl transfer. **(D)** Chemical representation of the two cofactors SAM and ProSeDMA.

Beyond its biochemical interest, SAMURI also provides a useful system for studying RNA-ligand interactions by computational approaches. This type of atomistic analysis is particularly valuable for RNA systems, where conformational heterogeneity, solvation, and ion binding are often tightly coupled determinants of structure and reactivity.^38,41^ Its relatively simple catalytic mechanism makes it well suited for examining cofactor conformational behavior in the active site, the determinants of cofactor stabilization, and the effects of mutations or cofactor modifications on reactivity.

In this work, we apply a multiscale^39,42–44^ computational enzymology framework^45,46^ to probe SAMURI’s catalytic mechanism at atomistic resolution. Molecular dynamics (MD) simulations were performed on the ribozyme-substrate-cofactor complex with either SAM or ProSeDMA, enabling a direct comparison of the two cofactors in the same structural context. Hybrid quantum mechanical/molecular mechanical (QM/MM) calculations were then used to explore the reaction coordinate, characterize the effect of mutations at the target adenosine A52 and evaluate the energy barriers. Free-energy calculations were also used to assess whether a divalent ion stabilizing the SAM cofactor is favorable. This combined approach corroborates the experimentally proposed S_N_2 -like mechanism, reveals key factors governing N3 selectivity and provides general principles for designing programmable RNA alkyltransferases for applications in chemical biology, RNA labeling and synthetic biology.

## Methods

### Molecular Dynamics simulations

#### System Preparation

Two main systems were prepared: the SAMURI ribozyme bound to either S-adenosylmethionine (SAM) or Propargylic Se-2,6-diaminopurin-ribosyl-selenomethionineamide (ProSeDMA). Starting structures were generated using VMD^47^ by superimposing the asymmetric units from the respective X-ray crystal structures (PDB codes 9FN2 for SAM and 9FN3 for ProSeDMA).^37^ Combining structural information from both subunits allowed localization of five Mg^2+^ ions in the SAM-bound structure and three in the ProSeDMA-bound structure. For each case, the simulated ribozyme was the subunit with the highest number of base pairs: the second subunit in 9FN2 and the first subunit in 9FN3. Two guanosine diphosphates originally present to improve crystal packing between subunits were removed. All crystallographically resolved water molecules within the Mg^2+^ hydration shells were retained. The SAMURI– cofactor complexes were parameterized with the ff99OL3 RNA force field^48,49^ and solvated in a truncated octahedron TIP4P-Ew water box.^50^ In all cases, ion counts ensured overall neutrality and reproduced a physiological 140 mM NaCl concentration, using monovalent ion parameters compatible with TIP4P-Ew and including 12–6–4 corrections. ^51,52^ For Mg^2+^ ions, the corrections of Panteva *et al.* were applied to improve the description of RNA–Mg^2+^ interactions.^53,54^

#### Cofactors and simulated systems

Classical molecular dynamics simulations were carried out for four cofactor states (Figure 2A): decarboxylated S-adenosylmethionine (SAM dcAdoMet), SAM, SAM + Mg^2+^, and ProSeDMA. QM/MM simulations were performed for all systems except SAM dcAdoMet and were further extended to the SAM + Mg^2+^ c^7^A52, ProSeDMA c^7^A52, and ProSeDMA c^1^A52 variants (Figures 3). Full system compositions are provided in Table S1 Supporting Information.

#### Parameterization of nonstandard residues

Three ligands and two modified nucleic acid residues were parametrized for use in the simulations. A common protocol was applied to all five species to ensure consistency with the AMBER force field family: each molecule was first geometry-optimized in Gaussian at the MP2/6-31G* level,^55^ followed by electrostatic potential calculations at the HF/6-31G* level. Partial atomic charges were then determined using the RESP procedure.^56^ Atom types and force-field parameters were assigned through Antechamber^57^ using AMBER atom types, so that the nucleic-acid-like fragments of the modified residues remained consistent with the standard AMBER OL3 residues^48^ and preserved compatibility with unmodified nucleotides. Selenium-specific bonded and nonbonded parameters were refined from sulfur analogies and MP2/6-31G*^55^ calculations on model compounds (TableS2 and FigureS1) (see Supporting Information for more details).

#### MD simulation protocol

All classical molecular dynamics (MD) simulations were performed with AMBER24^58^ using GPU-accelerated MD.^59,60^ Temperature was controlled with a Langevin thermostat^61^ and pressure with Monte Carlo barostat^62^ at 300 K and 1 atm. Long-range electrostatics were treated with particle mesh Ewald^63,64^ using a 12 Å real-space cutoff, and covalent bonds involving hydrogen were constrained with SHAKE.^65^ A 1 fs time step was used during equilibration. Systems were equilibrated through a multistep protocol including restrained minimization, solvent relaxation, heating, density equilibration, and progressive release of positional and mechanistic restraints consistent with previous works.^36,66^ Global positional restraints were initially applied to solute heavy atoms, together with additional mechanistically relevant restraints defined through the DISANG file. For ProSeDMA-containing systems, extra restraints were introduced to preserve the active-site stacking and hydrogen-bonding network during equilibration. Full restraint definitions and force constants are provided in the Supporting Information.

After equilibration, four independent 350 ns production trajectories were generated for each system in the NPT ensemble at 300 K and 1 atm. The first 100 ns of each trajectory were discarded for the analysis. Hydrogen mass repartitioning^67^ (HMR) was employed during production, enabling a 4 fs time step with SHAKE.^65^ Representative equilibrated structures were then selected for subsequent QM/MM simulations based on an in-line fitness score. Further details are given in the Supporting Information.

### 3D-RISM

Single point three-dimensional reference interaction site model (3D-RISM)^68,69^ calculations were performed on the selected asymmetric units stripped of solvent and ions including Mg^2+^. First, site-site solvent susceptibilities for a solution of 10 mM Mg^2+^, 140 mM Na^+^, and 160 mM Cl^−^ (32,768 grid points, 0.025 Å spacing) were determined from dielectrically consistent RISM (DRISM) with the rism1d program, as described in ref 70. The modified direct inversion of the iterative subspace (MDIIS) approach was used to iteratively solve the DRISM equation with PSE2 closure to a residual tolerance of 10^−12^ at 298 K and a dielectric constant of 78.497 for bulk water. A constant density approach was used, and the density of water was assumed to be 55.296 M using the SPC/E water model.^71^ Single point 3D-RISM was performed on a 200×200×200 Å^3^ grid with a 100×100×100 Å^3^ solvation box without solvent-solvent interaction cutoff. PSE1 closure was used with a residual tolerance of 10^−4^.

### Alchemical free energy simulations of Mg^2+^ binding

Absolute binding free energy (ABFE) simulations for Mg^2+^ were computed using AMBER24^58,60^ and analyzed using `FE-ToolKit`^72^ distributed within the AmberTools.^59^ Two systems were considered: (i) Mg^2+^ bound to the G30 phosphate in the SAM-bound ribozyme and (ii) a reference dinucleotide monophosphate system in which Mg^2+^ is bound to a phosphate group positioned between a guanine and a cytosine residue. The ribozyme system was extracted from the classical MD simulations described above, whereas the reference system was constructed separately at the same NaCl concentration of 140 mM.

The free-energy simulations were performed on three trials (Figure S2) using the GPU-accelerated free energy engine in AMBER^73^ that integrates advanced features described in detail elsewhere.^74,75^ Simulations were performed using an optimized alchemical transformation pathway with second-order smoothstep softcore potential,^76^ and 36-window optimized phase space overlap^77^ *λ* schedule, and ACES^78,79^ enhanced sampling approach (`gti_add_sc` flag set to 25). Within the ACES approach, Hamiltonian replica exchange^80–84^ attempts were made every 20 steps. A *λ*-dependent harmonic restraint between Mg^2+^ and the phosphate oxygen was applied during the transformation, and its contribution in the dummy state^85^ removed analytically.^86^ Production sampling was performed after rigorous end-state equilibration for 5 ns in the NPT ensemble. Free energies were estimated with MBAR^87^ using a variational method^88^ implemented within `FE-ToolKit` package^72^ distributed with AmberTools.^59^ Full details about the equilibration procedures, *λ* schedules, and supporting analyses are provided in the Supporting Information.

### *Ab initio* QM/MM simulations

All QM/MM simulations were conducted using recently developed interoperable software infrastructure for free energy simulations using generalized QM/MM + machine learning potentials in Amber.^89,90^ *Ab inito* QM/MM umbrella sampling simulations at the PBE0/631G* level^91^ were used to investigate methyl and propargyl transfer reactions with long-ranged electrostatic interactions treated rigorously with the ambient potential composite Ewald method.^92^

Starting configurations for *ab initio* QM/MM sampling were generated using two protocols depending on the cofactor. For SAM systems, windows were obtained sequentially using the third-order density-functional tight-binding (DFTB3) Hamiltonian and the 3OB3-1 parameter set,^93–96^ employing semiempirical Ewald electrostatics,^97^ with 3 ps of equilibration per window at this level, followed by 1 ps of equilibration at the PBE0 level.^91^ For ProSeDMA systems, windows were generated directly at the PBE0 level (as DFTB3 parameters are not available for Se) using 0.5 ps of equilibration per window followed by 1 ps of final equilibration. In all cases, the reported free-energy profiles are based on 30 ps of *ab initio* QM/MM sampling per window at the PBE0/6-31G* level,^91,98^ with long-range electrostatic interactions treated using the ambient potential composite Ewald method.^92^

All QM/MM simulations used 10 Å real space cutoffs, and were carried out with a 1 fs time step without use of SHAKE in the QM region. The QM-MM boundary was treated with the hydrogen link-atom approach.^99,100^

The reactive state is defined by SAM or ProSeDMA, with the positively charged sulfur or selenium center linked to a methyl or propargyl group, respectively, oriented toward the adenine 52. The product state corresponds to the adenine base methylated or propargylated at the N3 position. The reactions were described by a single reaction coordinate corresponding to alkyl transfer,

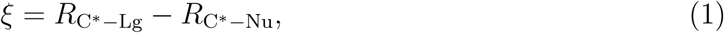

where *R*_C∗−Lg_ is the distance between the transferring carbon and the leaving group, and *R*_C∗−Nu_ is the distance between the transferring carbon and the nucleophile. Nu is m^3^A52 and proA52, Lg is SAH and SeDAH, respectively (Figure 3). Umbrella sampling was performed using 32 windows spanning *ξ*_SAM_ ∈ [−1.8, 2.4] Å and *ξ*_ProSeDMA_ ∈ [−1.6, 2.5] Å, with about 30 ps of sampling collected for each window. Free energy profiles were analyzed using a variational approach^101^ implemented within the `FE-ToolKit` package^72^ distributed with AmberTools.^59^

For SAM, the quantum region includes the adenine base of A52, the methyl group, the sulfur atom, and the entire tail of the cofactor, retaining the CH_2_ group adjacent to the sulfur center corresponding to carbon C5^′^ (cut between C5^′^ and C4^′^), for a total of atoms 37 atoms (Figure S3). In the SAM + Mg^2+^ conformation, the quantum part was expanded to comprise the Mg^2+^ ion with four coordinating water molecules, the phosphate group of G30 (including C3^′^, C5^′^, and bound hydrogens) and the SAM tail up to the ribose, giving a total of 60 atoms (Figure S3). Regarding the system with SAM as a cofactor interacting with Mg^2+^ and the c^7^A52 mutation, the QM region remains the same, except for a modification at base A52 that introduces one additional atom, resulting in a total of 61 atoms. For ProSeDMA, the quantum region includes the propargyl group, the selenium atom, the two CH_2_ groups directly bonded to selenium, the adenine base of A52, and the underlying guanine base G10 to capture *π*–*π* stacking with the propargyl triple bond, totaling 42 atoms (Figure S3). The definition of the QM part for the c^1^A and c^7^A variants was the same, with a change at base A52 leading to an additional atom (CH instead of N), with a total of 43 atoms. Detailed QM-region definitions, including the number of QM atoms for each system and the boundary choices, are provided in the Supporting Information.

#### *ab initio* DFT predictions of modified nucleobase p*K_a_*’s

A recent study made computational predictions of the p*K_a_* values at a number of positions for a wide range of modified nucleobases,^102^ including at the N3 position of adenine and the c^1^A and c^7^A modifications of specific interest to the present work. This study reported a predicted p*K_a_* value of 0.8 at the N3 position of adenine, which differed somewhat from the experimentally derived value of 1.5.^103^ Motivated by the original work,^102^ we set out to perform calculations that would provide a more precise estimate of the p*K_a_*shift at the N3 position of adenine due to c^1^A and c^7^A atomic mutants.

We performed p*K_a_* calculations on 15 modified nucleobases (Tables S3, S4 and Figure S4), closely related to adenine, using electronic structure and implicit solvation methods in Gaussian 16.^104^ Geometry optimization and frequency calculations were performed at the M06-2X/aug-cc-pVTZ level of theory^105,106^ with the SMD implicit solvation model.^107^ The p*K_a_* values were computed using a free energy approach based on the Gibbs free energy difference between protonated and deprotonated species in aqueous solution (Eq. 2), where the standard free energy of the proton in water (-270.297 kcal · mol^−1^) was used as in other work.^102^ Because only the relative p*K_a_* values (Δp*K_a_*) are considered, the contribution of the standard Gibbs free energy of the proton in water cancels and thus does not influence the calculated p*K_a_* shifts. The deprotonation Gibbs free energy for a species A is given as:

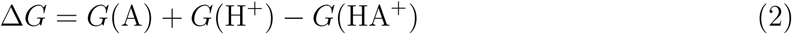

and the corresponding calculated p*K_a_* values were determined as:^102^

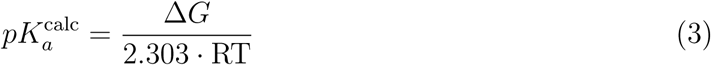

where R is the gas constant and T is temperature (298 K).

A linear regression model of the calculated p*K_a_* values was fit to the known experimental values^103,108–110^ to improve predictions of similar molecules with unknown p*K_a_* values. The resulting prediction model is given by

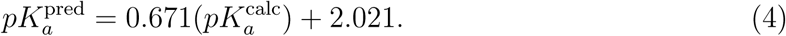

Additional details regarding the compounds used and validation tests are given in the Supporting Information.

## Results

### Conformational dynamics in solution indicate that the active ribozyme is a rare state

The conformational ensemble of SAMURI in the bound state was characterized in solution using molecular simulation. Specifically, the cofactors explicitly considered were SAM, a decarboxylated SAM variant (dcAdoMet), and ProSeDMA shown in Figure 2A. The available crystal structures capture a post-catalytic state with the alkylated target adenosine and the cofactor products bound in the active site.^31,37^ To generate models of the pre-reaction complexes, the product nucleoside and modified cofactor were replaced with their reactant counterparts, and then relaxed and equilibrated (see Computational Methods). Independent molecular dynamics simulations were then performed for each bound cofactor to generate conformational ensembles in solution (1 *µ*s cumulative sampling).

**Figure 2:**
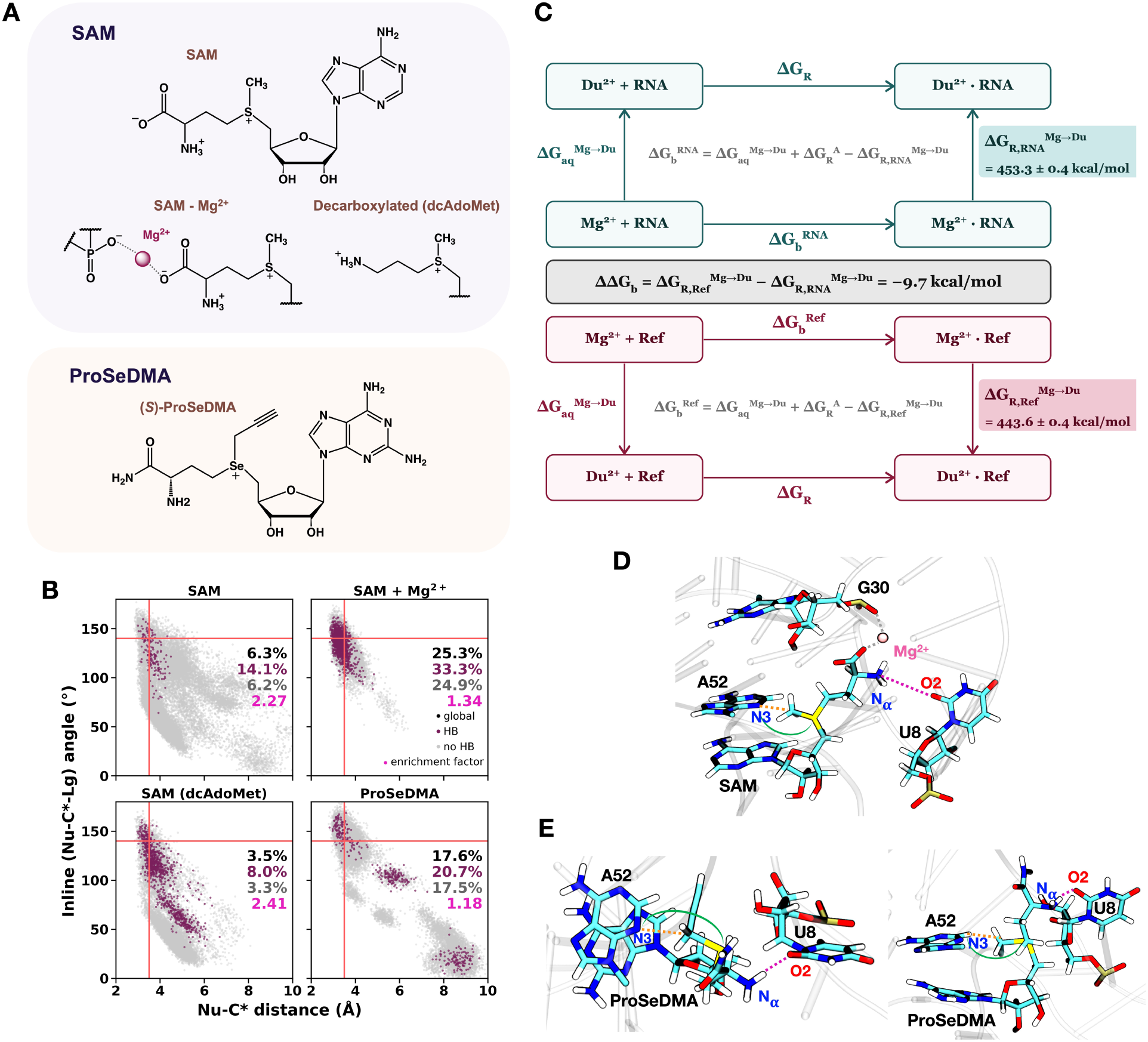
Active-site organization facilitates alkyl transfer. **(A)** Chemical structures of S-adenosylmethionine (SAM), SAM coordinated to Mg^2+^ (SAM-Mg^2+^), decarboxylated S-adenosylmethionine (dcAdoMet), and propargylic Se-2,6-diaminopurinribosylselenomethionineamide (ProSeDMA). **(B)** Distributions of inline attack angle (Nu-C^∗^-Lg) and distance (Nu-C^∗^), where the nucleophile (Nu) is A52:N3, C^∗^ is the electrophilic *α*-carbon of the cofactor, and the leaving group (Lg) is S (SAM, dcAdoMet) or Se (ProSeDMA). The fraction of reactive frames from four 250 ns replicas is shown in black (distance *<* 3.5 Å, angle *>* 140^◦^); frames with or without the *α*-amino–O2(U8) hydrogen bond are shown in burgundy and grey, respectively. **(C)**Thermodynamic cycle used to evaluate the relative binding free energy of the proposed Mg^2+^-binding site in SAMURI relative to a reference dinucleotide monophosphate, yielding ΔΔ*G_b_*. Here, Du^2+^ denotes a non-interacting dummy ion. 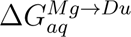 corresponds to the decoupling free energy of the ion in the bulk, 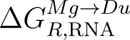 and 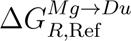 are the corresponding bound-state decoupling free energies, and Δ*G_R_* is the analytical restraint contribution. **(D)** Active site with SAM coordinated to the proposed Mg^2+^. **(E)** Active site with ProSeDMA (top and side views). In D and E, the nucleophilic attack distance (orange), attack angle (green), and the *α*-NH_2_–O2(U8) hydrogen bond (purple dashed lines) are indicated

All simulations showed stable ribozyme dynamics after ∼100 ns of relaxation from the crystal environment, preserving the overall fold and catalytic pocket (Figures S5–S8). The solvated ribozyme exhibited greater fluctuations than those estimated from crystallographic B factors, especially at peripheral, solvent-exposed helix termini and loop regions that were stabilized by packing contacts in the crystal (Figure S9). The catalytic core, on the other hand, remained comparatively more structured and less dynamic with root-mean-square deviation (RMSD) generally less than 2.0 Å.

In order for the S_N_2 alkylation reaction to occur, the nucleophile (A52:N3) needs to be in close proximity to the electrophilic *α* carbon of the cofactor (designated C^∗^) and form a nearly inline attack with the *α* carbon and leaving group (S in SAM, Se in ProSeDMA). Despite the structural stability of the active site, the relative positions of the reactive groups of the RNA and cofactor exhibit variation. Figure 2B shows the inline fitness distribution as a function of the inline attack angle and distance of the conformational ensembles in solution. The formation of a catalytically active state, characterized by a donor-acceptor distance of less than 3.5 Å and an inline attack angle greater than 140^◦^, only occurs 6.3% of the time for SAM and 17.6% of the time for ProSeDMA. This implies that the formation of catalytically active conformations for these cofactors is fairly rare (Figures S10 –S13).

### A predicted Mg^2+^ ion that anchors the tail and enhances inline fitness

Initial simulations of SAMURI were carried out without introducing additional divalent ions beyond those resolved in the crystallographic structure. Under these conditions for the SAM cofactor, Na^+^ ions were observed in the vicinity of the carboxylate terminus of the tail and a nearby phosphate (G30). Ribozyme active sites are often electrostatically strained^70^ and recruit a threshold occupancy of cationic charge from solvent.^111^ We thus employed 3D-RISM^68,112^ molecular solvation theory, which has been demonstrated to be successful in predicting functional metal ion binding sites in RNA,^70,113,114^ to explore a possible Mg^2+^ binding mode. Analysis revealed a pronounced Mg^2+^ density maximum between the SAM carboxylate and the pro-*S* _P_ oxygen of G30, supporting this region as a plausible binding site (Figure 2D and Figure S14). Another pronounced Mg^2+^ density maximum has been identified between C11 and U12 (Figure S15).

We next sought to gain more quantitative computational evidence for the feasibility of the Mg^2+^ binding site suggested by 3D-RISM using alchemical free energy simulations^74,75^ with enhanced sampling^77–79^ (see Computational Methods for details). The Mg^2+^ ion was initially positioned at the local maximum of the 3D-RISM density (Figure S14) and in direct coordination with both the carboxylate of the SAM cofactor and the pro-*S* _P_ oxygen of G30. Mg^2+^ ions are expected to bind fairly weakly to a single phosphate in, for example, a canonical RNA helix environment,^115,116^ whereas substantially stronger binding can emerge at specific RNA sites depending on the local structural and electrostatic context.^116,117^ The relative binding free energy, ΔΔ*G_b_*, of the Mg^2+^ ion in the proposed site in SAMURI with respect to an isolated dinucleotide monophosphate in solution was calculated to be -9.7 kcal · mol^−1^ using the thermodynamic cycle in Figure 2C (Figure S2). This indicates that the proposed Mg^2+^ binding site in SAMURI is predicted to be considerably more favorable than weak binding to the dinucleotide monophosphate reference. Taken together, this provides support that a Mg^2+^ ion could be partially occupied at this site for the SAM cofactor.

A notable outcome of the simulations is the effect of Mg^2+^ binding at this site, which has broader implications for ribozyme reactivity. The introduction of Mg^2+^ results in clear stabilization of both the global fold and the active site, with reduced local fluctuations and overall more stable behavior (Figures S5–S6, S9–S11). Remarkably, the overall inline fitness increased fourfold, from 6.3% in the absence of Mg^2+^ to 25.3% with Mg^2+^ bound (Figure 2B,D). This improvement arises primarily from reduced fluctuations in the cofactor tail, which are further enhanced by hydrogen bonding between the protonated *α* amine of the SAM cofactor and the O2 atom of U8. The broader implications of this interaction are discussed in the following subsection.

### An *α*-amine· · · U8–O2 hydrogen bond promotes catalytically competent conformations

Building on the experimental study,^37^ a key contact was identified between the cofactor *α*-amine and the O2 atom of U8, which is apparent in the crystal structure. In that work, deletion of A7, leaving U8 as the sole linker nucleotide, was sufficient to support folding and catalytic activity; subsequent substitution of U8 by cytidine was similarly well tolerated, whereas purine substitutions reduced activity, giving the trend U ≈ C *>* A *>* G. These observations point to a preference for pyrimidines and suggest that the O2 atom of U8 (present in both U and C) may contribute to catalysis as a hydrogen-bond acceptor, without being strictly required. Consistent with this interpretation, ProSeDAB and ProSeDMA, which retain the *α*-amine, display comparable experimental activity, whereas ProSeDMA– OH and ProSeDBA are substantially less active^37^ (see Figure S16).

To explore the role of U8:O2 in promoting inline fitness, simulations were performed for each of the cofactors shown in Figure 2A. Trajectories were clustered into non-overlapping sets according to the presence or absence of a hydrogen bond between the *α*-amine of the cofactors and U8:O2. Separate analysis of the inline fitness for both data sets enables the calculation of an enrichment factor, defined as the ratio of inline fitness values in the presence and absence of the hydrogen bond with U8:O2. In each case, the enrichment factors were all greater than unity, indicating that the presence of this hydrogen bond is correlated with enhanced inline fitness (Figure 2B). This effect is most pronounced in the least preorganized systems (SAM and dcAdoMet, enrichment factors 2.3-2.4), and more modest for the structured systems (SAM+Mg^2+^ and ProSeDMA, enrichment factors 1.2-1.3) that had greater overall inline fitness. In all cases explored in the current work, the origin of enhanced inline fitness is correlated with the introduction of order in the cofactor tail, and thus depends on the broader hydrogen bond interaction network of the cofactor with key residues in the active site (Table S5), of which U8:O2 plays a prominent role (Figures S10– S13).

### Conformational preorganization and electrostatic tuning shape the free-energy barrier for alkyl transfer

The observed rate for a pseudo first-order enzymatic reaction can be modeled as *k_obs_* = *f_react_* · *k_int_*, where *f_react_* is the fraction of the dynamical ensemble of the enzyme in a catalytically competent reactive conformation (henceforth referred to as the “active state”), and *k_int_* is the intrinsic rate departing from the enzyme in its active state. In general, the active state fraction is a function of reaction conditions^118^ (e.g., pH, ion environment, and temperature), whereas the intrinsic rate (sometimes referred to as the cleavage rate for RNA enzymes^119^) derives from the chemical steps of the reaction and depends only on temperature under the assumption that the mechanism itself is invariant to the other state variables. For a given enzyme, both *f_react_*and *k_int_* are expected to vary with different substrates. Assuming simple transition state theory (unit transmission coefficient^120^), the intrinsic rate can be modeled as *k_int_* = (*k_B_T/h*)*e*^−Δ*G*‡^*^/^*^(*k*^*^BT^* ^)^, where Δ*G*^‡^ is the activation free energy, *k_B_* and *h* are the Boltzmann and Planck constants, respectively, and *T* is the Kelvin temperature. In the case of acid-base catalysis, experimental activity-pH profiles can often be used to extract kinetic parameters that can be used to estimate *f_react_* and *k_int_*, although these may not always be uniquely determined due to the principle of kinetic ambiguity. ^118^ The SAMURI does not employ acid-base catalysis, and only the experimentally observed rate constant, *k_obs_*, is available.^31,37^ Nonetheless, based on the molecular simulation data presented here, we can make an estimate of *f_react_* for SAM and ProSeDMA based on the active state populations presented above. Using the percentages of inline fitness shown in Figure 2B, the active state fractions can be approximated as 0.063 and 0.176 for SAM and ProSeDMA, respectively. The corresponding experimentally observed rates^31,37^ are 4×10^−4^ min^−1^ and 1.49×10^−1^ min^−1^, leading to estimated intrinsic rates of 6.35 × 10^−3^ min^−1^ and 8.47 × 10^−1^ min^−1^. From transition state theory, the activation free energy values for SAM and ProSeDMA, derived from the experimental rate and calculated active state fraction estimates, are 22.9 and 20.0 kcal · mol^−1^ for SAM and ProSeDMA, respectively. These values are used to compare and validate the activation free energy values derived from *ab initio* QM/MM simulations described below. The alkyl-transfer reactions considered here for SAM and ProSeDMA are shown schematically in Figure 3A,B. In the SAM system, the reaction corresponds to the transfer of a methyl group from sulfur to the N3 of A52, whereas in the ProSeDMA system, a propargyl group is transferred from selenium to the same nucleophilic center. In both cases, the reaction proceeds through an S_N_2 mechanism, with cleavage of the S/Se–C^∗^ bond and formation of the C^∗^–N3 bond.

**Figure 3:**
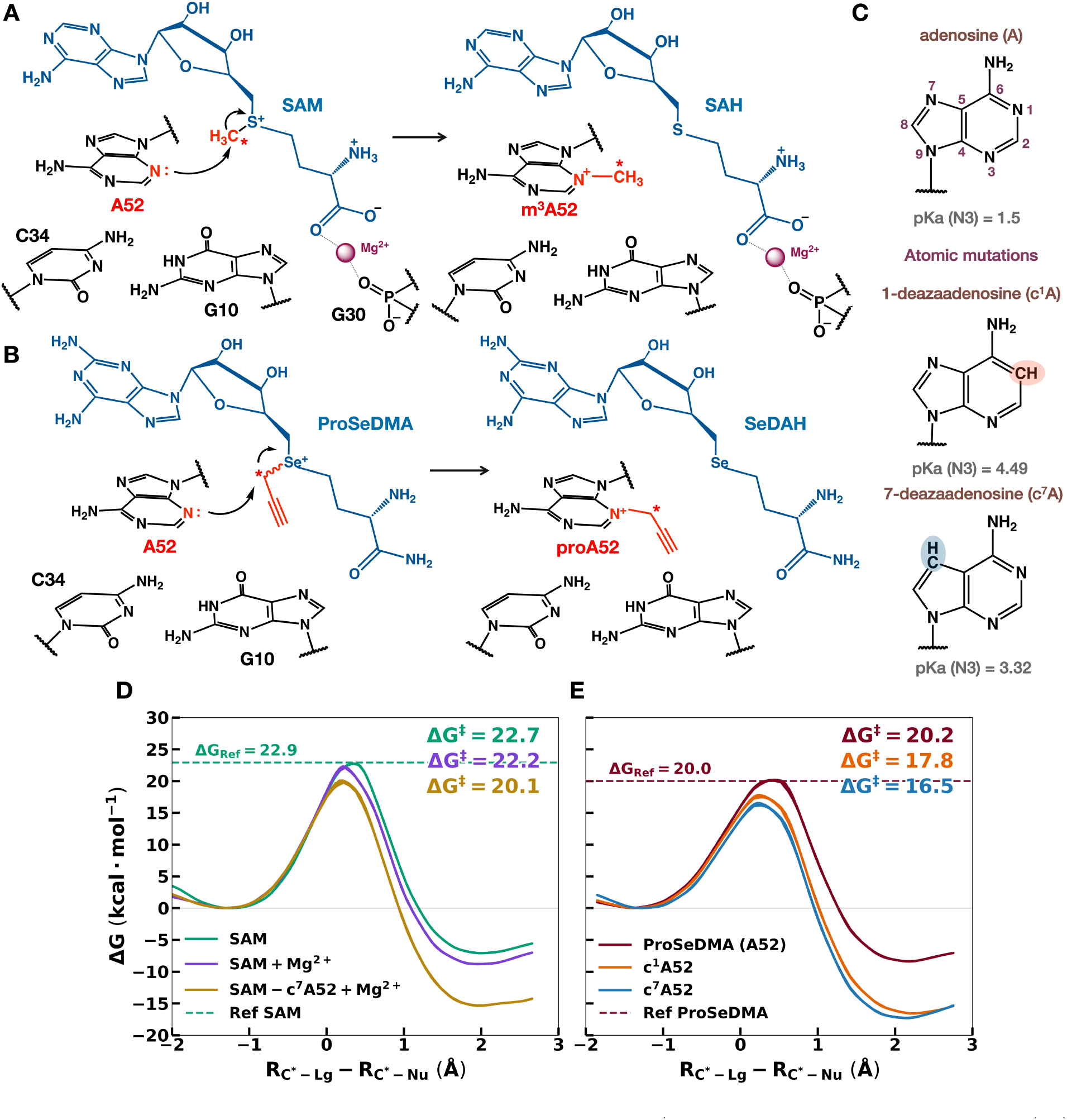
Alkyl-group transfer reactions and QM/MM free-energy profiles. **(A)** ChemDraw representation of the methyl transfer from SAM to N3 of A52. **(B)** ChemDraw representation of the propargyl transfer from ProSeDMA to N3 of A52. **(C)** p*K_a_*(N3) values for adenosine (A) from ref.,^103^ 1-deazaadenosine (c^1^A) and 7-deazaadenosine (c^7^A) estimated from shifts relative to A calculated (Tables S3, S4). **(D)** *ab initio* QM/MM free-energy profiles projected onto the reaction coordinate *R*_C∗−Lg_ −*R*_C∗−Nu_, with activation free energies (Δ*G*^‡^). Profiles were obtained from umbrella sampling (32 windows, ∼30 ps per window). Left: SAM, SAM+Mg^2+^, SAM+c^7^A+Mg^2+^. Right: comparison for A (ProSeDMA), c^1^A and c^7^A substrates. In both cases, the experimental values are represented in dashed lines.

The reaction progress was described by a single distance-difference coordinate, *R*_C∗−Lg_ −*R*_C∗−Nu_, where C^∗^ is the reacting *α* carbon, Lg is the leaving group (S in SAM and Se in ProSeDMA), and Nu is the nucleophile (N3 of A52). Free energy profiles^101^ from *ab initio* QM/MM simulations with full long-ranged Ewald electrostatics^92^ are shown in Figure 3D,E. The activation free energy for SAM is Δ*G*^‡^ = 22.7 kcal · mol^−1^ in the absence of Mg^2+^, and slightly lower, Δ*G*^‡^ = 22.2 kcal · mol^−1^ in the presence of Mg^2+^ (Figure 3D). The lowering of the activation free energy in the presence of the bound Mg^2+^ results from preferential electrostatic interactions with the ion in the product state relative to the reactant state.

Specifically, in going from the reactant state to the product, a formal +1 charge migrates from the leaving group to the nucleophile position, which is located farther away from the Mg^2+^. This results in a modest lowering of the reaction free energy relative to the reaction in the absence of the Mg^2+^, and in accordance with expected linear free energy relations,^121–124^ also leads to an earlier transition state and a slight lowering of the forward activation free energy barrier. The calculated activation free energy values are in close agreement (0.2-0.7 kcal · mol^−1^) with the reference activation free energy of 22.9 kcal · mol^−1^.

The activation free energy for ProSeDMA is Δ*G*^‡^ = 20.2 kcal · mol^−1^, in excellent agreement with the reference value of 20.0 kcal · mol^−1^. The calculated and reference activation barriers for ProSeDMA are 2.0–2.9 kcal · mol^−1^ lower than those for SAM, consistent with selenium’s larger size and greater polarizability relative to sulfur, which enhance its ability to act as a leaving group.

### QM/MM simulations aid in the interpretation of atomic mutagenesis experiments and enable new predictions

Fundamental experimental studies of nucleic acid enzymes have provided a wealth of insight into their structural organization and catalytic strategies.^12,125–127,127^ In the work introducing the SAMURI,^31^ variants of the ribozyme at the A52 position were explored in conjunction with the ProSeDMA cofactor. In particular, 1-deazaadenosine (c^1^A) and 7-deazaadenosine (c^7^A) atomic mutants were examined, and showed similar reactivity with ProSeDMA (although the rates were not specifically measured). This deaza substitution replaces an electron withdrawing endocyclic nitrogen in the ring with a more electron donating CH group, which is expected to alter the electronic structure and possibly the activity. We thus calculated QM/MM free energy profiles to predict the effects of these modifications on the activation barriers and intrinsic rate.

The activation free energy of c^1^A is 17.8 kcal · mol^−1^, whereas c^7^A is 16.5 kcal · mol^−1^ (Figure 3E). Hence, these atomic mutants have barriers lower than the unmodified SAMURI by 2.4 and 3.7 kcal · mol^−1^, respectively. These trends are consistent with the expected correlation with the microscopic p*K_a_* values at the N3 position shown in Figure 3C.^102,103^

Indeed, both c^1^A and c^7^A analogs have upshifted N3 p*K_a_*values (by 2.4-4.1 units) relative to the unmodified adenine (Figure 3C). The resulting electron donation into the aromatic ring leads to increased nucleophilicity at the N3 position and preferentially stabilizes the product state making the reaction more exergonic. In accordance with linear free energy relationships,^121–124^ this lowers the forward barrier and shifts the transition state toward reactants (producing an “earlier” transition state). In addition to these electronic effects, there are some differences observed in the hydrogen bond network between the c^1^A and c^7^A simulations. Specifically, for c^1^A, the N7 position receives a hydrogen bond from the HO2^′^ of C34, whereas in the c^7^A simulation this interaction is not possible. Rather, in the c^7^A simulation, the N6 exocyclic amine donates a hydrogen bond to the O6 of G36 which forms a netwrok with the N4 of C11 and N1 of A52.

Given the results for the c^1^A and c^7^A atomic mutants for ProSeDMA, we set out to predict a modification in the SAM cofactor that would be practical and enhance the ribozyme reactivity. In the SAMURI study,^31^ it was reported that the electrospray ionization (ESI)-MS trace of the alkylated c^1^A sample contained a depurinated fragment not seen in the c^7^A sample. This behavior can be rationalized by the classical depurination mechanism, in which protonation at N7 generates a cationic purine that weakens the N-glycosidic bond and promotes cleavage (Figure S17). By contrast, c^7^A prevents protonation at N7 and consequently blocks access to the depurination pathway.^31^ To avoid the unwanted side reaction with c^1^A, we focus on the c^7^A atomic mutant in the presence of a bound Mg^2+^ ion to explore the degree to which electronic effects might further enhance the reaction rate with the SAM cofactor. Since this mutation is expected to increase the p*K_a_* at N3, it was of interest to test whether the geometric effects induced by Mg^2+^ and the electronic effects associated with the c^7^A substitution could have a multiplicative effect on the rate. Analogously to ProSeDMA cofactor, the calculated free energy profile for the c^7^A atomic mutant for the SAM cofactor leads to stabilization of the product state, lowering of the activation barrier by 2.6 and 2.1 kcal · mol^−1^ with respect to the unmodified cofactor in the absence and presence of bound Mg^2+^, respectively. These results suggest that atomic mutants of A52 that increase the nucleophilicity at N3, together with structural elements that anchor the cofactor tail may be fruitful in enhancing SAMURI reactivity.

## Discussion

The SAMURI is an in vitro-selected ribozyme that binds SAM and ProSeDMA cofactors to catalyze site-specific RNA alkylation, including the transfer of a propargyl group used as a tag for azide-alkyne click chemistry in RNA labeling.^31,37^ The SAMURI does not appear to engage in sophisticated acid or base chemistry to achieve catalysis. Rather, the SAMURI structure binds the cofactor and positions the *α* carbon (C^∗^) in sufficiently close proximity to the alkylation site (N3 of A52) so that reactive conformations can be sampled. Simulations suggest that these reactive conformations, collectively referred to as the “active state”, are fairly rare, and the introduction of structural elements that stabilize these conformations would further enhance the reactivity. In particular, results suggest that the low fraction of reactive frames observed for SAM is consistent with a more disordered cofactor tail in comparison with the ProSeDMA that appears somewhat better adapted to the active state ensemble.

This scenario is similar to that of the self-alkylating ribozyme^128,129^ (SAR), which catalyzes the alkylation of the N7 atom of a guanine by loosely binding a polyether ligand with a reactive terminal epoxide and positioning it to undergo nucleophilic attack and ring opening. Alternatively, this mechanistic strategy contrasts with that reported for the methyltransferase ribozyme^130,131^ (MTR1), in which the reactive arrangement is much more strongly biased toward an inline geometry.^36,132^ The MTR1 active site is substantially more preorganized for chemistry than in SAMURI, using a protonated cytosine residue as an acid catalyst and an elaborate hydrogen bond network to position the reactive elements and stabilize the nucleobase leaving group.

Hence, building upon the MTR1 example, strategies to improve catalytic efficiency in SAMURI would involve introducing interactions that further stabilize the active state, or introducing environmental or chemical modifications to enhance the reactivity. Simulations predict that one such example of the former might be achieved by the binding of a Mg^2+^ ion interacting with the SAM carboxylate, which is consistent with structural motifs observed in other SAM-binding RNAs.^25,133–135^ As summarized in Figure S18, divalent ions in several RNA systems contribute directly to the organization of the cofactor tail by bridging the carboxylate to the RNA scaffold or to nearby nucleobases. For example, in the SAM-V riboswitch, Mg^2+^ connects the SAM carboxylate to a backbone phosphate, ^133^ whereas in the SAM-I riboswitch a divalent ion directly coordinates the cofactor carboxylate while also interacting with a neighboring guanine.^134^ A related principle is also observed in the SAM-dependent methyltransferase ribozyme, where a divalent metal coordinates the SAM-like tail together with neighboring nucleobase heteroatoms.^25^ Taken together, these examples support the idea that divalent ions can locally stabilize charge and reinforce the architecture of SAM-binding pockets through a range of coordination modes. Results presented here from 3D-RISM and alchemical free energy simulations provide support for the plausible existence of a similar Mg^2+^ binding site; however, the absence of such an observed metal in the crystal structure suggests that if such a binding site exists, it is likely fairly weak and at most only partially occupied. Nonetheless, the simulation results support the supposition that if a stronger divalent metal ion binding site could be engineered, it is likely to result in more efficient catalysis with the SAM cofactor.

The simulation results also implicate the importance of the *α*-amine· · · U8–O2 hydrogen bond in stabilizing the active state. This interaction appears to function primarily as a local organizing contact within the SAMURI active site, and correlates with the enrichment of reactive conformations, consistent with a role of biasing the conformational ensemble toward catalytically competent states. This effect is most evident in the least preorganized SAM systems, whereas its relative contribution is attenuated in SAM+Mg^2+^ and ProSeDMA, where the cofactor is more conformationally constrained. These results support a model in which the *α*-amine· · · U8–O2 interaction contributes to catalytic organization as one of several cooperative local interactions.^37^

The free energy profiles reveal trends in the intrinsic reactivity of the chemical steps that can be rationalized through consideration of electronic effects. The activation barriers are lower for the ProSeDMA cofactor than for SAM owing to the electronic properties of Se that enable it to be an enhanced leaving group relative to S. In addition, the propargyl group of ProSeDMA is electron withdrawing, which can better stabilize partial positive charge on the electrophillic C^∗^ compared to SAM. This electron withdrawing property is analogous to the enhanced reactivity of MTR1 in the presence of *O*^6^-alkylguanine moieties carrying aromatic substituents on C^∗^. Atomic mutants that increase electron donation into the nucleobase aromatic ring raise the microscopic p*K_a_*and enhance the nucleophilicity of the N3 position, leading to lower barriers and a faster intrinsic rate. Other isofunctional substitutions on A52 that tune the p*K_a_* of N3 could lead to further enhancement.

Alternatively, there may be ways of altering the active site electrostatic environment that lead to p*K_a_* tuning at the A52:N3 position to enhance its reactivity. For example, the twister ribozyme^136–138^ is a small self-cleaving RNA that uses the N3 position of a conserved active site adenine (A1) to act as a general acid catalyst.^46,139,140^ However, the intrinsic p*K_a_* of the N3 position of adenine is quite low (estimated to be 1.5^103^), and hence, if unshifted, would not be available in its protonated form required to act as an acid catalyst at near-neutral pH. The twister ribozyme active site, however, has positioned two phosphates to dually coordinate the N6 exocyclic amine of A1, anchoring it in a position to act as a general acid by transferring a proton to the O5^′^ leaving group of the scissile phosphate. At the same time, these two anionic residues serve to shift the p*K_a_* of A1 at the N3 position by approximately 3.5 units.^46,140^ It is possible that engineering similar interactions with the N6 exocyclic amine of A52 in SAMURI could further shift the p*K_a_* and enhance the nucleophilicity at the N3 position.

## Conclusion

By integrating molecular dynamics, 3D-RISM solvation analysis, alchemical free-energy calculations, quantum p*K_a_* predictions, and *ab initio* QM/MM free-energy simulations, we delineate the conformational and electronic determinants of SAMURI catalysis. Together, these results establish that the origin of catalysis derives from a synergy between conformational preorganization and electronic tuning, in which rare reactive conformations and intrinsic chemical reactivity jointly determine activity. This mechanistic framework provides general design principles for engineering programmable RNA catalysts for site-specific chemical modification, advancing their application in chemical biology and therapeutics.

## Acknowledgement

The authors are grateful for financial support provided by the National Institutes of Health (No. GM062248 to D.M.Y.). E.M. acknowledges support from the National Science Foundation Graduate Research Fellowship under Grant No. 2233066. Authors thank A.S.H. for the valuable discussions and sharp comments. Computational resources were provided by the Office of Advanced Research Computing (OARC) at Rutgers, The State University of New Jersey; the Advanced Cyberinfrastructure Coordination Ecosystem: Services & Support (ACCESS) program, which is supported by National Science Foundation grants #2138259, #2138286, #2138307, #2137603, and #2138296 (supercomputer Expanse at SDSC through allocation CHE190067); and the Texas Advanced Computing Center (TACC) at the University of Texas at Austin, URL: http://www.tacc.utexas.edu (supercomputer Frontera through allocation CHE20002).

## References

(1) Cech, T. R. The Chemistry of Self-Splicing RNA and RNA Enzymes. Science 1987, 236, 1532–1539.

(2) Kruger, K.; Grabowski, P.; Zaug, A.; Sands, J.; Gottschling, D.; Cech, T. Self-splicing RNA: autoexcision and autocyclisation of the ribosomal RNA intervening sequence of *Tetrahymena*. Cell 1982, 31, 147–157.

(3) Guerrier-Takada, C.; Gardiner, K.; Maresh, T. The RNA moiety of ribonuclease P is the catalytic subunit of the enzyme. Cell 1983, 35, 849–857.

(4) Doherty, E. A.; Doudna, J. A. Ribozyme Structures and Mechanisms. Annu. Rev. Biophys. Biomol. Struct. 2001, 30, 457–475.

(5) Gilbert, W. The RNA World. Nature 1986, 319, 618.

(6) E., O. L. Prebiotic Chemistry and the Origin of the RNA World. Crit. Rev. Biochem. Mol. Biol. 2004, 39, 99–123.

(7) Hanna, R.; Doudna, J. A. Metal ions in ribozyme folding and catalysis. Curr. Opin. Chem. Biol. 2000, 4, 166–170.

(8) Scott, W. G. Ribozymes. Curr. Opin. Struct. Biol. 2007, 17, 280–286.

(9) Wilson, C.; Szostak, J. In vitro evolution of a self-alkylating ribozyme. Nature 1995, 374, 777–782.

(10) Yokobayashi, Y. High-throughput analysis and engineering of ribozymes and deoxyribozymes by sequencing. Acc. Chem. Res. 2020, 53, 2903–2912.

(11) Lassila, J. K.; Zalatan, J. G.; Herschlag, D. Biological phosphoryl-transfer reactions: Understanding mechanism and catalysis. Annu. Rev. Biochem. 2011, 80, 669–702.

(12) Lilley, D. M. J. Classification of the nucleolytic ribozymes based upon catalytic mechanism. F1000 Res. 2019, 8, 1462.

(13) Bevilacqua, P. C.; Harris, M. E.; Piccirilli, J. A.; Gaines, C.; Ganguly, A.; Kostenbader, K.; Ekesan, Ş.; York, D. M. An Ontology for Facilitating Discussion of Catalytic Strategies of RNA-Cleaving Enzymes. ACS Chem. Biol. 2019, 14, 1068–1076.

(14) Zhang, B.; Cech, T. R. Peptide bond formation by in vitro selected ribozymes. Nature 1997, 390, 96–100.

(15) Doudna, J. A.; Cech, T. R. The chemical repertoire of natural ribozymes. Nature 2002, 418, 222–228.

(16) Lilley, D. M. J. The origins of RNA catalysis in ribozymes. Trends Biochem. Sci. 2003, 28, 495–501.

(17) Fedor, M. J.; Williamson, J. R. The catalytic diversity of RNAs. Nat. Rev. Mol. Cell Biol. 2005, 6, 399–412.

(18) Robertson, D. L.; Joyce, G. F. Selection in vitro of an RNA enzyme that specifically cleaves singlestranded DNA. Nature 1990, 344, 467–468.

(19) Johnston, W. K.; Unrau, P. J.; Lawrence, M. S.; Glasner, M. E.; Bartel, D. P. RNA-Catalyzed RNA Polymerization: Accurate and General RNA-Templated Primer Extension. Science 2001, 292, 1319–1325.

(20) Bartel, D. P.; Szostak, J. W. Isolation of new ribozymes from a large pool of random sequences. Science 1993, 261, 1411–1418.

(21) Tarasow, T. M.; Tarasow, S. L.; Eaton, B. E. RNA-catalysed carbon–carbon bond formation. Nature 1997, 389, 54–57.

(22) Fusz, S.; Eisenfuhr, A.; Srivatsan, S. G.; Heckel, A.; Famulok, M. A Ribozyme for the Aldol Reaction. Chem. Biol. 2005, 12, 941–950.

(23) Sengle, G.; Eisenfuhr, A.; Arora, P. S.; Nowick, J. S.; Famulok, M. Novel RNA catalysts for the Michael reaction. Chem. Biol. 2001, 8, 459–473.

(24) Scheitl, C. P. M.; Ghaem Maghami, M.; Lenz, A.-K.; Höbartner, C. Site-specific RNA methylation by a methyltransferase ribozyme. Nature 2020, 587, 663–667.

(25) Jiang, H.; Gao, Y.; Zhang, L.; Chen, D.; Gan, J.; Murchie, A. I. H. The identification and characterization of a selected SAM-dependent methyltransferase ribozyme that is present in natural sequences. Nat Catal 2021, 4, 872–881.

(26) Scheitl, C. P.; Dorinova, E.; Christopher, S.; Walunj, M. B.; Okuda, T.; Sednev, M. V.; Höbartner, C. An Unexpected Adenosine-Alkylating Ribozyme Emerged by Target Site Relocation during in Vitro Selection. J. Am. Chem. Soc. 2025, 147, 44225–44235.

(27) Dorinova, E.; Walunj, M. B.; Höbartner, C. Alkyltransferase Ribozyme for SiteSpecific N4-Cytidine Alkylation. Angew. Chem. Int. Ed. 2026, 6447137.

(28) Batey, R. T. Recognition of *S* -adenosylmethionine by riboswitches. WIREsRNA 2011, 2, 299–311.

(29) El Yacoubi, B.; Bailly, M.; de Crécy-Lagard, V. Biosynthesis and Function of Posttranscriptional Modifications of Transfer RNAs. Annu. Rev. Genet. 2012, 46, 69–95.

(30) Sharma, S.; Watzinger, P.; Kötter, P.; Entian, K.-D. Identification of a novel methyltransferase, Bmt2, responsible for the N-1-methyl-adenosine base modification of 25S rRNA in *Saccharomyces cerevisiae*. Nucleic Acids Res. 2013, 41, 5428–43.

(31) Okuda, T.; Lenz, A.-K.; Seitz, F.; Vogel, J.; Höbartner, C. A SAM analogue-utilizing ribozyme for site-specific RNA alkylation in living cells. Nat. Chem. 2023, 15, 1523–1531.

(32) Maghami, M.; Scheitl, C. P. M.; Höbartner, C. Direct *in Vitro* Selection of *Trans*Acting Ribozymes for Posttranscriptional, Site-Specific, and Covalent Fluorescent Labeling of RNA. J. Am. Chem. Soc. 2019, 141, 19546–19549.

(33) Scheitl, C. P. M.; Okuda, T.; Adelmann, J.; Höbartner, C. Ribozyme-Catalyzed Late-Stage Functionalization and Fluorogenic Labeling of RNA. Angew. Chem. Int. Ed. 2023, 62, 202305463.

(34) Fontecave, M.; Atta, M.; Mulliez, E. S-adenosylmethionine: nothing goes to waste. Trends Biochem. Sci. 2004, 29, 243–249.

(35) Du, J.; Mahcene, B.; Martynov, V.; Frezza, E.; Vasnier, C.; Ponchon, L.; Coelho, D.; Bonhomme, F.; Braud, E.; Etheve-Quelquejeu, M. et al. Investigation of a squaramide motif as a bioisostere of the amino-acid group of S-adenosyl-L-methionine and its functional impact on RNA methylation. Commun. Chem. 2025, 8, 244.

(36) McCarthy, E.; Ekesan, Ş.; Giese, T. J.; Wilson, T. J.; Deng, J.; Huang, L.; Lilley, D. M. J.; York, D. M. Catalytic mechanism and pH dependence of a methyltransferase ribozyme (MTR1) from computational enzymology. Nucleic Acids Res. 2023, 51, 4508–4518.

(37) Chen, H.-A.; Okuda, T.; Lenz, A.-K.; Scheitl, C. P.; Schindelin, H.; Höbartner, C. Structure and catalytic activity of the SAM-utilizing ribozyme SAMURI. Nat. Chem. Biol. 2025,

(38) Auffinger, P.; Westhof, E. RNA solvation: A molecular dynamics simulation perspective. Biopolymers 2001, 56, 266–274.

(39) Palermo, G.; Cavalli, A.; Klein, M. L.; Alfonso-Prieto, M.; Dal Peraro, M.; De Vivo, M. Catalytic metal ions and enzymatic processing of DNA and RNA. Acc. Chem. Res. 2015, 48, 220–228.

(40) Sgrignani, J.; Magistrato, A. The Structural Role of Mg^2+^ Ions in a Class I RNA Polymerase Ribozyme: A Molecular Simulation Study. J. Phys. Chem. B 2012, 116, 2259–2268.

(41) Auffinger, P.; Louise-May, S.; Westhof, E. Molecular Dynamics Simulations of Solvated Yeast tRNA *^PHE^*. Biophys. J. 1999, 76, 50–64.

(42) Nam, K.; Gao, J.; York, D. M. In Multiscale Simulation Methods for Nanomaterials; Ross, R. B., Mohanty, S. S., Eds.; John Wiley & Sons, Inc., 2008; pp 201–218.

(43) Christensen, A. S.; Kubař, T.; Cui, Q.; Elstner, M. Semiempirical Quantum Mechanical Methods for Noncovalent Interactions for Chemical and Biochemical Applications. Chem. Rev. 2016, 116, 5301–5337, PMID: 27074247.

(44) Barros, E. P.; Ries, B.; Böselt, L.; Champion, C.; Riniker, S. Recent developments in multiscale free energy simulations. Curr. Opin. Struct. Biol. 2022, 72, 55–62.

(45) Panteva, M. T.; Dissanayake, T.; Chen, H.; Radak, B. K.; Kuechler, E. R.; Giambaşu, G. M.; Lee, T.-S.; York, D. M. In Methods in Enzymology; Chen, S.-J., Burke-Aguero, D. H., Eds.; Elsevier, 2015; Vol. 553; Chapter 14.

(46) Gaines, C. S.; Giese, T. J.; York, D. M. Cleaning Up Mechanistic Debris Generated by Twister Ribozymes Using Computational RNA Enzymology. ACS Catal. 2019, 9, 5803–5815.

(47) Humphrey, W.; Dalke, A.; Schulten, K. VMD: Visual Molecular Dynamics. J. Mol. Graphics 1996, 14, 33–38.

(48) Zgarbová, M.; Otyepka, M.; Šponer, J.; Mládek, A.; Banáš, P.; Cheatham III, T. E.; Jurečka, P. Refinement of the Cornell et al. nucleic acids force field based on reference quantum chemical calculations of glycosidic torsion profiles. J. Chem. Theory Comput. 2011, 7, 2886–2902.

(49) Pérez, A.; Marchán, I.; Svozil, D.; Sponer, J.; Cheatham III, T. E.; Laughton, C. A.; Orozco, M. Refinement of the AMBER force field for nucleic acids: Improving the description of *α/γ* conformers. Biophys. J. 2007, 92, 3817–3829.

(50) Horn, H. W.; Swope, W. C.; Pitera, J. W.; Madura, J. D.; Dick, T. J.; Hura, G. L.; Head-Gordon, T. Development of an improved four-site water model for biomolecular simulations: TIP4P-Ew. J. Chem. Phys. 2004, 120, 9665–9678.

(51) Li, P.; Song, L. F.; Merz Jr, K. M. Systematic Parameterization of Monovalent Ions Employing the Nonbonded Model. J. Chem. Theory Comput. 2015, 11, 1645–57.

(52) Sengupta, A.; Li, Z.; Song, L. F.; Li, P.; Merz Jr., K. M. Parameterization of Monovalent Ions for the OPC3, OPC, TIP3P-FB, and TIP4P-FB Water Models. J. Chem. Inf. Model. 2021, 61, 869–880.

(53) Panteva, M. T.; Giambaşu, G. M.; York, D. M. Comparison of structural, thermodynamic, kinetic and mass transport properties of Mg^2+^ ion models commonly used in biomolecular simulations. J. Comput. Chem. 2015, 36, 970–982.

(54) Panteva, M. T.; Giambasu, G. M.; York, D. M. Force Field for Mg^2+^, Mn^2+^, Zn^2+^, and Cd^2+^ Ions that have Balanced Interactions with Nucleic Acids. J. Phys. Chem. B 2015, 119, 15460–15470.

(55) Møller, C.; Plesset, M. S. Note on an approximation treatment for many-electron systems. Phys. Rev. 1934, 46, 618–622.

(56) Schauperl, M.; Nerenberg, P. S.; Jang, H.; Wang, L.-P.; Bayly, C. I.; Mobley, D. L.; Gilson, M. K. Non-bonded force field model with advanced restrained electrostatic potential charges (RESP2). Commun. Chem. 2020, 3, 44.

(57) Wang, J.; Wang, W.; Kollman, P. A.; Case, D. A. Automatic atom type and bond type perception in molecular mechanical calculations. J. Mol. Graph. Model. 2006, 25, 247–260.

(58) Case, D.; Aktulga, H.; Belfon, K.; Ben-Shalom, I.; Berryman, J.; Brozell, S.; Cerutti, D.; Cheatham, I., T.E.; Cisneros, G.; Cruzeiro, V., et al. Amber 2024 Reference Manual. University of California, San Francisco, 2024.

(59) Case, D. A.; Aktulga, H. M.; Belfon, K.; Cerutti, D. S.; Cisneros, G. A.; Cruzeiro, V. W. D.; Forouzesh, N.; Giese, T. J.; Götz, A. W.; Gohlke, H., et al. AmberTools. J. Chem. Inf. Model. 2023, 63, 6183–6191.

(60) Case, D. A.; Cerutti, D. S.; Cruzeiro, V. W. D.; Darden, T. A.; Duke, R. E.; Ghazimirsaeed, M.; Giambaşu, G. M.; Giese, T. J.; Götz, A. W.; Harris, J. A., et al. Recent Developments in Amber Biomolecular Simulations. J. Chem. Inf. Model. 2025, 65, 7835–7843.

(61) Loncharich, R. J.; Brooks, B. R.; Pastor, R. W. Langevin dynamics of peptides: the frictional dependence of isomerization rates of N-acetylalanyl-N’-methylamide. Biopolymers 1992, 32, 523–535.

(62) Åqvist, J.; Wennerström, P.; Nervall, M.; Bjelic, S.; Brandsdal, B. O. Molecular dynamics simulations of water and biomolecules with a Monte Carlo constant pressure algorithm. Chem. Phys. Lett. 2004, 384, 288–294.

(63) Darden, T.; York, D.; Pedersen, L. Particle mesh Ewald: An N log(N) method for Ewald sums in large systems. J. Chem. Phys. 1993, 98, 10089–10092.

(64) Essmann, U.; Perera, L.; Berkowitz, M. L.; Darden, T.; Lee, H.; Pedersen, L. G. A smooth particle mesh Ewald method. J. Chem. Phys. 1995, 103, 8577–8593.

(65) Ryckaert, J. P.; Ciccotti, G.; Berendsen, H. J. C. Numerical Integration of the Cartesian Equations of Motion of a System with Constraints: Molecular Dynamics of nAlkanes. J. Comput. Phys. 1977, 23, 327–341.

(66) Ekesan, Ş.; York, D. M. Dynamical ensemble of the active state and transition state mimic for the RNA-cleaving 8-17 DNAzyme in solution. Nucleic Acids Res. 2019, 47, 10282–10295.

(67) Hopkins, C. W.; Le Grand, S.; Walker, R. C.; Roitberg, A. E. Long-Time-Step Molecular Dynamics through Hydrogen Mass Repartitioning. J. Chem. Theory Comput. 2015, 11, 1864–1874.

(68) Luchko, T.; Gusarov, S.; Roe, D. R.; Simmerling, C.; Case, D. A.; Tuszynski, J.; Kovalenko, A. Three-dimensional molecular theory of solvation coupled with molecular dynamics in AMBER. J. Chem. Theory Comput. 2010, 6, 607–624.

(69) Genheden, S.; Luchko, T.; Gusarov, S.; Kovalenko, A.; Ryde, U. An MM/3D-RISM Approach for Ligand Binding Affinities. J. Phys. Chem. B 2010, 114, 8505–8516.

(70) Ekesan, Ş.; McCarthy, E.; Case, D. A.; York, D. M. RNA Electrostatics: How Ribozymes Engineer Active Sites to Enable Catalysis. J. Phys. Chem. B 2022, 126, 5982–5990.

(71) Berendsen, H. J. C.; Grigera, J. R.; Straatsma, T. P. The Missing Term in Effective Pair Potentials. J. Phys. Chem. 1987, 91, 6269–6271.

(72) Giese, T. J.; Snyder, R.; Piskulich, Z.; Barletta, G. P.; Zhang, S.; McCarthy, E.; Ekesan, Ş.; York, D. M. FE-ToolKit: A Versatile Software Suite for Analysis of HighDimensional Free Energy Surfaces and Alchemical Free Energy Networks. J. Chem. Inf. Model. 2025, 65, 5273–5279.

(73) Lee, T.-S.; Cerutti, D. S.; Mermelstein, D.; Lin, C.; LeGrand, S.; Giese, T. J.; Roitberg, A.; Case, D. A.; Walker, R. C.; York, D. M. GPU-Accelerated Molecular Dynamics and Free Energy Methods in Amber18: Performance Enhancements and New Features. J. Chem. Inf. Model. 2018, 58, 2043–2050.

(74) Lee, T.-S.; Allen, B. K.; Giese, T. J.; Guo, Z.; Li, P.; Lin, C.; Jr., T. D. M.; Pearlman, D. A.; Radak, B. K.; Tao, Y., et al. Alchemical Binding Free Energy Calculations in AMBER20: Advances and Best Practices for Drug Discovery. J. Chem. Inf. Model. 2020, 60, 5595–5623.

(75) York, D. M. Modern Alchemical Free Energy Methods for Drug Discovery Explained. ACS Phys. Chem. Au 2023, 3, 478–491.

(76) Tsai, H.-C.; Lee, T.-S.; Ganguly, A.; Giese, T. J.; Ebert, M. C.; Labute, P.; Merz Jr, K. M.; York, D. M. AMBER Free Energy Tools: A New Framework for the Design of Optimized Alchemical Transformation Pathways. J. Chem. Theory Comput. 2023, 19, 640–658.

(77) Zhang, S.; Giese, T. J.; Lee, T.-S.; York, D. M. Alchemical Enhanced Sampling with Optimized Phase Space Overlap. J. Chem. Theory Comput. 2024, 20, 3935–3953.

(78) Lee, T.-S.; Tsai, H.-C.; Ganguly, A.; York, D. M. ACES: Optimized Alchemically Enhanced Sampling. J. Chem. Theory Comput. 2023, 19, 472–487.

(79) Tsai, H.; Xu, J.; Guo, Z.; Yi, Y.; Tian, C.; Que, X.; Giese, T.; Lee, T.-S.; York, D. M.; Ganguly, A. et al. Improvements in Precision of Relative Binding Free Energy Calculations Afforded by the Alchemical Enhanced Sampling (ACES) Approach. J. Chem. Inf. Model. 2024, 64, 7046–7055.

(80) Sindhikara, D.; Meng, Y.; Roitberg, A. E. Exchange frequency in replica exchange molecular dynamics. J. Chem. Phys. 2008, 128, 24103.

(81) Hritz, J.; Oostenbrink, C. Hamiltonian replica exchange molecular dynamics using soft-core interactions. J. Chem. Phys. 2008, 128, 144121.

(82) Yang, M.; Huang, J.; MacKerell, Jr., A. D. Enhanced Conformational Sampling Using Replica Exchange with Concurrent Solute Scaling and Hamiltonian Biasing Realized in One Dimension. J. Chem. Theory Comput. 2015, 11, 2855–2867.

(83) Jiang, W.; Roux, B. Free energy perturbation Hamiltonian replica-exchange molecular dynamics (FEP/H-REMD) for absolute ligand binding free energy calculations. J. Chem. Theory Comput. 2010, 6, 2559–2565.

(84) Arrar, M.; de Oliveira, C. A. F.; Fajer, M.; Sinko, W.; McCammon, J. A. w-REXAMD: A Hamiltonian replica exchange approach to improve free energy calculations for systems with kinetically trapped conformations. J. Chem. Theory Comput. 2013, 9, 18– 23.

(85) Fleck, M.; Wieder, M.; Boresch, S. Dummy Atoms in Alchemical Free Energy Calculations. J. Chem. Theory Comput. 2021, 17, 4403–4419.

(86) Boresch, S. On Analytical Corrections for Restraints in Absolute Binding Free Energy Calculations. J. Chem. Inf. Model. 2024, 64, 3605–3609.

(87) Shirts, M. R.; Chodera, J. D. Statistically optimal analysis of samples from multiple equilibrium states. J. Chem. Phys. 2008, 129, 124105.

(88) Giese, T. J.; York, D. M. Variational Method for Networkwide Analysis of Relative Ligand Binding Free Energies with Loop Closure and Experimental Constraints. J. Chem. Theory Comput. 2021, 17, 1326–1336.

(89) Giese, T. J.; Zeng, J.; Lerew, L.; McCarthy, E.; Tao, Y.; Ekesan, Ş.; York, D. M. Software Infrastructure for Next-Generation QM/MM-ΔMLP Force Fields. J. Phys. Chem. B 2024, 128, 6257–6271.

(90) Tao, Y.; Giese, T. J.; Şölen Ekesan; Zeng, J.; Aradi, B.; Hourahine, B.; Aktulga, H. M.; Götz, A. W.; Merz, Jr, K. M.; York, D. M. Amber free energy tools: Interoperable software for free energy simulations using generalized quantum mechanical/molecular mechanical and machine learning potentials. J. Chem. Phys. 2024, 160, 224104.

(91) Adamo, C.; Cossi, M.; Barone, V. An accurate density functional method for the study of magnetic properties: the PBE0 model. J. Mol. Struct. (Theochem) 1999, 193, 145–157.

(92) Giese, T. J.; York, D. M. Ambient-Potential Composite Ewald Method for *ab Initio* Quantum Mechanical/Molecular Mechanical Molecular Dynamics Simulation. J. Chem. Theory Comput. 2016, 12, 2611–2632.

(93) Elstner, M.; Porezag, D.; Jungnickel, G.; Elsner, J.; Haugk, M.; Frauenheim, T.; Suhai, S.; Seifert, G. Self-consistent-charge density-functional tight-binding method for simulations of complex materials properties. Phys. Rev. B 1998, 58, 7260–7268.

(94) Cui, Q.; Elstner, M.; Kaxiras, E.; Frauenheim, T.; Karplus, M. A QM/MM Implementation of the Self-Consistent Charge Density Functional Tight Binding (SCC-DFTB) Method. J. Phys. Chem. B 2001, 105, 569–585.

(95) Gaus, M.; Cui, Q.; Elstner, M. DFTB3: Extension of the seld-consistent-charge density-functional tight-binding method (SCC-DFTB). J. Chem. Theory Comput. 2011, 7, 931–948.

(96) Gaus, M.; Goez, A.; Elstner, M. Parametrization and Benchmark of DFTB3 for Organic Molecules. J. Chem. Theory Comput. 2013, 9, 338–354.

(97) Nam, K.; Gao, J.; York, D. M. An efficient linear-scaling Ewald method for longrange electrostatic interactions in combined QM/MM calculations. J. Chem. Theory Comput. 2005, 1, 2–13.

(98) Hariharan, P. C.; Pople, J. A. The Influence of Polarization Functions on Molecular Orbital Hydrogenation Energies. Theor. Chim. Acta 1973, 28, 213–222.

(99) Warshel, A.; Levitt, M. Theoretical studies of enzymic reactions: Dielectric, electrostatic and steric stabilization of the carbonium ion in the reaction of lysozyme. J. Mol. Biol. 1976, 103, 227–249.

(100) Singh, U. C.; Kollman, P. A. A combined *ab initio* quantum mechanical and molecular mechanical method for carrying out simulations on complex molecular systems: Applications to the CH_3_Cl+Cl^−^ exchange reaction and gas phase protonation of polyethers. J. Comput. Chem. 1986, 7, 718–730.

(101) Giese, T. J.; Ekesan, Ş.; York, D. M. Extension of the Variational Free Energy Profile and Multistate Bennett Acceptance Ratio Methods for High-Dimensional Potential of Mean Force Profile Analysis. J. Phys. Chem. A 2021, 125, 4216–4232.

(102) Mlotkowski, A. J.; Schlegel, H. B.; Chow, C. S. Calculated pK*_a_* Values for a Series of Aza- and Deaza-Modified Nucleobases. J. Phys. Chem. A 2023, 127, 3526–3534.

(103) Kapinos, L. E.; Operschall, B. P.; Larsen, E.; Sigel, H. Understanding the acid-base properties of adenosine: the intrinsic basicities of N1, N3 and N7. Chem. Eur. J. 2011, 17, 8156–8164.

(104) Frisch, M. J.; Trucks, G. W.; Schlegel, H. B.; Scuseria, G. E.; Robb, M. A.; Cheeseman, J. R.; Scalmani, G.; Barone, V.; Petersson, G. A.; Nakatsuji, H., et al. Gaussian 16, Revision C.01. Gaussian, Inc.: Wallingford, CT, 2016.

(105) Zhao, Y.; Truhlar, D. G. The M06 suite of density functionals for main group thermochemistry, thermochemical kinetics, noncovalent interactions, excited states, and transition elements: two new functionals and systematic testing of four M06-class functionals and 12 other functionals. Theor. Chem. Acc. 2008, 120, 215–241.

(106) Kendall, R. A.; Dunning, Jr., T. H.; Harrison, R. J. Electron affinities of the first-row atoms revisited. Systematic basis sets and wave functions. J. Chem. Phys. 1992, 96, 6796–6806.

(107) Marenich, A. V.; Cramer, C. J.; Truhlar, D. G. Universal Solvation Model Based on Solute Electron Density and on a Continuum Model of the Solvent Defined by the Bulk Dielectric Constant and Atomic Surface Tensions. J. Phys. Chem. B 2009, 113, 6378–6396.

(108) Bereiter, R.; Himmelstoß, M.; Renard, E.; Mairhofer, E.; Egger, M.; Breuker, K.; Kreutz, C.; Ennifar, E.; Micura, R. Impact of 3-deazapurine nucleobases on RNA properties. Nucleic Acids Res. 2021, 49, 4281–4293.

(109) Wierzchowski, J.; Wielgus-Kutrowska, B.; Shugar, D. Fluorescence emission properties of 8-azapurines and their nucleosides, and application to the kinetics of the reverse synthetic reaction of purine nucleoside phosphorylase. Biochim. Biophys. Acta 1996, 1290, 9–17.

(110) Martin, D. M. G.; Reese, C. B. Some aspects of the chemistry of N(1)and N(6)dimethylallyl derivatives of adenosine and adenine. J. Chem. Soc. C 1968, 0, 1731–1738.

(111) Lee, T.-S.; Giambaşu, G. M.; Sosa, C. P.; Martick, M.; Scott, W. G.; York, D. M. Threshold Occupancy and Specific Cation Binding Modes in the Hammerhead Ribozyme Active Site are Required for Active Conformation. J. Mol. Biol. 2009, 388, 195–206.

(112) Kovalenko, A.; Truong, T. N. Thermochemistry of solvation: A self-consistent threedimensional reference interaction site model approach. J. Chem. Phys. 2000, 113, 7458–7470.

(113) Giambaşu, G. M.; Case, D. A.; York, D. M. Predicting Site-Binding Modes of Ions and Water to Nucleic Acids Using Molecular Solvation Theory. J. Am. Chem. Soc. 2019, 141, 2435–2445.

(114) Ganguly, A.; Weissman, B. P.; Giese, T. J.; Li, N.-S.; Hoshika, S.; Rao, S.; Benner, S. A.; Piccirilli, J. A.; York, D. M. Confluence of theory and experiment reveals the catalytic mechanism of the Varkud satellite ribozyme. Nat. Chem. 2020, 12, 193–201.

(115) Sigel, R. K. O.; Sigel, H. A stability concept for metal ion coordination to singlestranded nucleic acids and affinities of individual sites. Acc. Chem. Res. 2010, 43, 974–984.

(116) Cunha, R. A.; Bussi, G. Unraveling Mg(2+)-RNA binding with atomistic molecular dynamics. RNA 2017, 23, 628–638.

(117) Leonarski, F.; D’Ascenzo, L.; Auffinger, P. Mg^2+^ ions: do they bind to nucleobase nitrogens? Nucleic Acids Res. 2017, 45, 987–1004.

(118) Bevilacqua, P. C. Mechanistic considerations for general acid-base catalysis by RNA: Revisiting the mechanism of the hairpin ribozyme. Biochemistry 2003, 42, 2259–2265.

(119) Wilson, T. J.; Liu, Y.; Li, N. S.; Dai, Q.; Piccirilli, J. A.; Lilley, D. M. Comparison of the structures and mechanisms of the pistol and hammerhead ribozymes. J. Am. Chem. Soc. 2019, 141, 7865–7875.

(120) Garcia-Viloca, M.; Gao, J.; Karplus, M.; Truhlar, D. G. How enzymes work: Analysis by modern rate theory and computer simulations. Science 2004, 303, 186–195.

(121) Jencks, W. P. A Primer for the Bema Hapothle. An Empirical Approach to the Characterization of Changing Transition-State Structures. Chem. Rev. 1985, 85, 511–527.

(122) Williams, A. Effective charge and Leffler’s index as mechanistic tools for reactions in solution. Acc. Chem. Res. 1984, 17, 425–430.

(123) Huang, M.; York, D. M. Linear free energy relationships in RNA transesterification: theoretical models to aid experimental interpretations. Phys. Chem. Chem. Phys. 2014, 16, 15846–15855.

(124) Chen, H.; Giese, T. J.; Huang, M.; Wong, K.-Y.; Harris, M. E.; York, D. M. Mechanistic Insights into RNA Transphosphorylation from Kinetic Isotope Effects and Linear Free Energy Relationships of Model Reactions. Chem. Eur. J. 2014, 20, 14336–14343.

(125) Micura, R.; Höbartner, C. Fundamental studies of functional nucleic acids: aptamers, riboswitches, ribozymes and DNAzymes. Chem. Soc. Rev. 2020, 49, 7331–7353.

(126) Wilson, T. J.; Lilley, D. M. J. The potential versatility of RNA catalysis. Wiley Interdiscip. Rev. RNA 2021, 12, 1651.

(127) Lilley, D. M. J.; Huang, L. RNA catalysis moving towards metabolic reactions: progress with ribozyme catalyzed alkyl transfer. Trends Biochem. Sci. 2025, 50, 417–424.

(128) McDonald, R. I.; Guilinger, J. P.; Mukherji, S.; Curtis, E. A.; Lee, W. I.; Liu, D. R. Electrophilic activity-based RNA probes reveal a self-alkylating RNA for RNA labeling. Nat. Chem. Biol. 2014, 10, 1049–54.

(129) Krochmal, D.; Shao, Y.; Li, N.-S.; DasGupta, S.; Shelke, S. A.; Koirala, D.; Piccirilli, J. A. Structural basis for substrate binding and catalysis by a self- alkylating ribozyme. Nat. Chem. Biol. 2022, 18, 376–384.

(130) Scheitl, C. P. M.; Mieczkowski, M.; Schindelin, H.; Höbartner, C. Structure and mechanism of the methyltransferase ribozyme MTR1. Nat. Chem. Biol. 2022, 18, 547–555.

(131) Deng, J.; Wilson, T. J.; Wang, J.; Peng, X.; Li, M.; Lin, X.; Liao, W.; Lilley, D. M. J.; Huang, L. Structure and mechanism of a methyltransferase ribozyme. Nat. Chem. Biol. 2022, 18, 556–564.

(132) Wilson, T. J.; McCarthy, E.; Ekesan, Ş.; Giese, T. J.; Li, N.-S.; Huang, L.; Piccirilli, J. A.; York, D. M.; Lilley, D. M. J. The Role of General Acid Catalysis in the Mechanism of an Alkyl Transferase Ribozyme. ACS Catal. 2024, 14, 15294–15305.

(133) Huang, L.; Lilley, D. M. Structure and ligand binding of the SAM-V riboswitch. Nucleic Acids Res. 2018, 46, 6869–6879.

(134) Huang, L.; Wang, J.; Lilley, D. A critical base pair in k-turns determines the conformational class adopted, and correlates with biological function. Nucleic Acids Res. 2016, 44, 5390–5398.

(135) Lu, C.; Smith, A. M.; Fuchs, R. T.; Ding, F.; Rajashankar, K.; Henkin, T. M.; Ke, A. Crystal structures of the SAM-III/SMK riboswitch reveal the SAM-dependent translation inhibition mechanism. Nat. Struct. Mol. Biol. 2008, 15, 1076–1083.

(136) Liu, Y.; Wilson, T. J.; McPhee, S. A.; Lilley, D. M. J. Crystal structure and mechanistic investigation of the twister ribozyme. Nat. Chem. Biol. 2014, 10, 739–744.

(137) Ren, A.; Košutić, M.; Rajashankar, K. R.; Frener, M.; Santner, T.; Westhof, E.; Micura, R.; Patel, D. J. In-line alignment and Mg2+ coordination at the cleavage site of the env22 twister ribozyme. Nat. Commun. 2014, 5, 5534–5544.

(138) Ren, A.; Vusurovic, N.; Gebetsberger, J.; Gao, P.; Juen, M.; Kreutz, C.; Micura, R.; Patel, D. Pistol Ribozyme Adopts a Pseudoknot Fold Facilitating Site-specific In-line Cleavage. Nat. Chem. Biol. 2016, 12, 702–708.

(139) Wilson, T. J.; Liu, Y.; Domnick, C.; Kath-Schorr, S.; Lilley, D. M. J. The Novel Chemical Mechanism of the Twister Ribozyme. J. Am. Chem. Soc. 2016, 138, 6151–6162.

(140) Gaines, C. S.; York, D. M. Ribozyme Catalysis with a Twist: Active State of the Twister Ribozyme in Solution Predicted from Molecular Simulation. J. Am. Chem. Soc. 2016, 138, 3058–3065.

