## Supporting Information for "Origins of reactivity in SAM-utilizing ribozyme SAMURI-catalyzed RNA alkylation"

### Detailed Computational Methods

#### Detailed composition of simulated systems

**Table S1: Details of all systems builds including modifications and solvent components**

| Cofactor | Modification | Water | Na | Cl | Mg |
|---|---|---|---|---|---|
| MD and QM/MM | | | | | |
| SAM | | 10,874 | 72 | 28 | 6 |
| SAM | $+Mg^{2+}$ | 11,075 | 70 | 28 | 7 |
| SAM | dcAdoMet | 11,139 | 72 | 29 | 6 |
| ProSeDMA | | 12,173 | 79 | 31 | 4 |
| QM/MM | | | | | |
| SAM | $+Mg^{2+}$,$c^7$A52 | 11,075 | 70 | 28 | 7 |
| ProSeDMA | $c^7$A52 | 12,173 | 79 | 31 | 4 |
| ProSeDMA | $c^1$A52 | 12,173 | 79 | 31 | 4 |

Cofactors included S-adenosylmethionine (SAM), decarboxylated S-adenosylmethionine (dcAdoMet), and propargylic Se-2,6-diaminopurinribosyl-selenomethionineamide (ProSeDMA).$+Mg^{2+}$ indicates the addition of a $Mg^{2+}$ ion coordinating the cofactor tail.

#### Non-standard residue parametrization

ProSeDMA was parameterized using a fragment-based procedure within the AMBER framework.[1] In this protocol, the cofactor was divided into three components: the diaminopurine nucleobase, the ribose moiety and the selenium-containing side chain. Charges for the diaminopurine fragment were first derived by RESP fitting of electrostatic potentials computed with Gaussian.[2,3] Charges for the sugar fragment were taken directly from the corresponding ribose fragment available in AMBER, using the variant consistent with the desired protonation state for the O3′ hydroxyl group. The selenium-containing tail was parameterized separately as an independent methyl capped fragment. Its geometry was optimized at the MP2/6-31G* level of theory prior to electrostatic-potential calculations and RESP charge derivation. This fragment includes the Se-containing propargyl group together with the rest of the side chain attached to the ribose, so that the final charge distribution remained consistent over the whole substituent. The three fragments were then assembled into a single cofactor description. In addition, the atom type of C5′ was reverted from CT to simplify the parameterization.

An initial parameter set for the assembled cofactor was then generated. Because ProSeDMA contains selenium, this automatically generated parameter set required additional manual refinement. Initial selenium parameters were adapted from sulfur analogies and these terms were subsequently refined using MP2/6-31G* optimizations[4] performed on tripropylselenium and tripropylsulfonium model compounds. Comparison of these two reference systems was used to adjust the selenium-specific force-field terms and improve the description of the Se-containing part of the cofactor (Figure S1 and Table S2).

#### ProSeDMA parameters

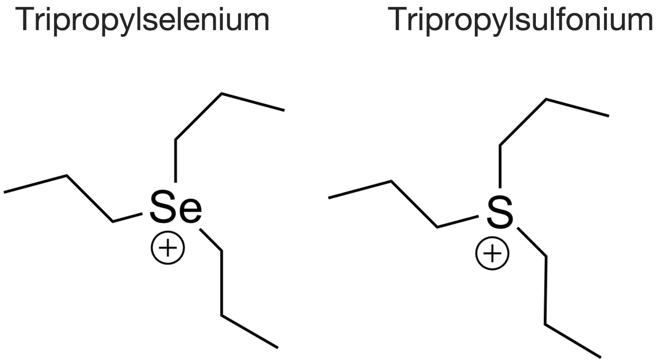


Figure S1: Representation of the tripropylselenonium and tripropylsulfonium model compounds used to derive the force-field parameters, highlighting the differences induced by replacing sulfur with selenium

**Table S2: Comparison of force-field parameters for selenium (Se) and sulfur (S)**

| Parameter | Se | S |
|---|---|---|
| Mass (amu) | 78.871 | 32.060 |
| Bond length C3–X (Å) | 1.952 | 1.831 |
| Angle C3–X–C3 (deg) | 99.493 | 96.120 |
| Angle C1–C3–X (deg) | 110.989 | 106.350 |
| Angle C3–C3–X (deg) | 111.530 | 110.120 |
| Angle H1–C3–X (deg) | 107.920 | 107.920 |
| $\sigma$ (Å) | 2.180 | 2.000 |
| $\epsilon$ ($\mathrm{kcal \cdot mol^{-1}}$) | 0.220 | 0.250 |
| $\alpha$ ($\mathrm{Å^3}$) | 3.77 | 3.00 |

##### Atomic mutagenesis of the target adenosine

Because 1-deazaadenosine ($c^1A$) and 7-deazaadenosine ($c^7A$) are not standard AMBER residues, they were parameterized separately prior to system construction. Partial charges were obtained using the same Gaussian/RESP procedure as for the other nonstandard cofactors. Atom types were assigned from AMBER by selecting the closest possible analogies to those of native adenine, in order to reproduce the parent adenosine base as closely as possible.

Using the ProSeDMA-containing system, mutated variants were then introduced at the A52 base, where alkyl transfer takes place. For the $c^1A$ and $c^7A$ systems, the initial structures were generated directly from the equilibrated umbrella-sampling windows of the native adenosine (A) system. The mutations were introduced from the corresponding A structures in each of the 32 equilibrated umbrella-sampling windows to ensure consistency across all starting configurations.

#### MD simulation protocol

The system was equilibrated through a multistep protocol including solvent relaxation, gradual heating, density stabilization, and staged release of positional and mechanistic restraints. Initially, global positional restraints were applied to all solute heavy atoms with a force constant of $k = 50\ \mathrm{kcal \cdot mol^{-1} \cdot Å^{-2}}$.

In parallel, relevant restraints were applied using the same force constant. These included: (i) semi-harmonic restraints preserving the crystallographic $Mg^{2+}$ crystal-packing ions, (ii) a restraint on the $Mg^{2+}$–carboxylate contact (target distance 3.6 Å), (iii) a restraint on the N$\alpha$H···O2(U8) hydrogen bond (target distance 2.63 Å), (iv) a semi-harmonic upper-bound restraint on the nucleophile–electrophile distance (boundary at 3.0 Å), (v) a semi-harmonic angular restraint enforcing an in-line attack geometry (lower bound 170°). For ProSeDMA system, additional restraints were introduced: (vi) to maintain the propargyl group of ProSeDMA in a $\pi$-stacking arrangement over the G10 base, three flat-bottom distance restraints were applied between the propargyl moiety and G10, keeping distances within the 3.2–4.2 Å range and preventing separation from the guanine surface; (vii) three flat-bottom distance restraints were applied to preserve key interactions between G9, U37, and the cofactor, namely between G9(O6) and U37(O2′), G9(N1) and U37(O2), and ProSeDMA(N6) and U37(O4). These restraints were defined such that no penalty was applied below 3.2 Å, while a harmonic potential was applied beyond this distance. They were necessary to maintain the structural integrity of the active site, as their absence led to disruption of these interactions during the simulations.

This was followed by a procedure of progressive restraint reduction, as described below. An initial solvent minimization was performed while maintaining the positional restraints ($k$ = 50), followed by three short NPT simulations at 300 K and 1 atm (5 ps, 5 ps, and 190 ps) to stabilize the box dimensions and density. The system was then re-minimized and heated under constant volume from 0 to 300 K over 600 ps, held at 300 K for 1 ns, and further equilibrated for 5 ns in the NPT ensemble. During all these preparation, heating, and early equilibration stages, both positional (`ntr`) and relevant restraints (`DISANG`) were maintained at $k = 50\ \mathrm{kcal \cdot mol^{-1} \cdot \mathring{A}^{-2}}$. Subsequently, the global positional restraints on the solute were progressively reduced through consecutive equilibration stages (25, 10, 5, and 2 $\mathrm{kcal \cdot mol^{-1} \cdot \mathring{A}^{-2}}$, while all restraints remained fixed at $k = 50$. Once positional restraints were fully removed (`ntr = 0`), an extended equilibration phase was carried out in which only the relevant restraints were retained. A first 50 ns segment was performed with $k = 40$, followed by successive 10 ns segments in which the force constants associated with the defined restraints were progressively reduced to 30, 20, and 10 $\text{kcal} \cdot \text{mol}^{-1} \cdot \text{Å}^{-2}$. This gradual release allowed controlled relaxation of the hydrogen bond and in-line attack restraints while avoiding abrupt structural perturbations. For the SAM-containing systems, two $Mg^{2+}$ ions from the crystallographic environment were retained even during production. These ions were relatively unstable during equilibration (drift close to the active site); therefore, weak restraints ($k \approx 40$ $\text{kcal} \cdot \text{mol}^{-1} \cdot \text{Å}^{-2}$) were maintained on both crystallographic $Mg^{2+}$ ions throughout production to preserve their crystallographic positions. In contrast, for the ProSeDMA systems, the crystallographic packing $Mg^{2+}$ ion remained structurally stable during equilibration. Consequently, the restraints applied to the crystallographic $Mg^{2+}$ ions were progressively reduced following the staged protocol described above, allowing gradual relaxation. However, the restraints between residues G9 and U37, as well as between ProSeDMA and U37, were maintained with a force constant of 30 $\text{kcal} \cdot \text{mol}^{-1}$.

From these classical simulations, representative equilibrated structures were selected as starting points for subsequent QM/MM investigations of the adenine N3-alkylation reaction based on a catalytic fitness score.

#### Analysis of catalytic fitness

Catalytic fitness of the simulated ensembles was evaluated by monitoring the interaction between the nucleophile and electrophile. Two descriptors were used: (i) the distance between the alkyl carbon of the transferable group (methyl or propargyl) and the nucleophilic N3 atom of adenine, and (ii) the angle defined by the donor heteroatom (S or Se), the alkyl carbon, and N3. For each trajectory frame, a normalized geometric score was computed as

$$\text{score} = \frac{\text{value} - \text{cutoff}}{\text{best} - \text{cutoff}}, \tag{1}$$

$$\text{combined score} = \sqrt{\frac{(\text{angle score})^2 + (\text{distance score})^2}{2}}. \tag{2}$$

For SAM-like and ProSeDMA-like cofactors best distance was set to 3.0 Å with a cutoff of 3.5 Å and the optimal angle was set to 170° with a cutoff of 140°. Frames below cutoff were assigned a score of zero. The resulting values ranged from 0 to 1 providing a direct measure of proximity to a catalytically relevant alignment.

### Binding free energy calculations

#### Equilibration of the real state

After a first minimization, an equilibration phase of about 1 ns in the NPT ensemble is performed to allow adjustment of the simulation box volume, the temperature control is ensured by a Langevin thermostat[5] with a friction coefficient of 2 $ps^{-1}$, whereas the pressure is regulated with Monte Carlo barostat.[6] Once stabilized, a simulation in the NVT ensemble is following during 500 ps. The main equilibration phase of the real state consists in molecular dynamics of about 2 ns. A heating and cooling cycle is then applied taking the system from 300 K to 600 K, maintaining this elevated temperature and then returning it to 300. This aims to improve conformational diversity within the replicas. In this process of equilibration of the real state, the time step used is 1 fs, and the cutoff for the long range interactions is 10 Å.

#### Alchemical transformation

Simulations were performed using an optimized alchemical transformation pathway with second-order smoothstep softcore potential and 36-windows optimized phase space overlap $\lambda$ schedule.[7] For the reference system: $\lambda \in$ [0.000000, 0.100340, 0.182850, 0.224850 0.260250, 0.290780, 0.316950, 0.340330, 0.357760, 0.369870, 0.380700, 0.392860, 0.409240, 0.428010, 0.446350, 0.465110, 0.483390, 0.499220, 0.513990, 0.529750, 0.546590, 0.563260, 0.578520,

0.593130, 0.609250, 0.628600, 0.649050, 0.665850, 0.681370, 0.700150, 0.727260, 0.754670, 0.783840, 0.820050, 0.899540, 1.000000]. For the RNA system: $\lambda \in$ [0.000000, 0.100510, 0.188890, 0.229220, 0.260910, 0.283890, 0.304900, 0.328100, 0.351030, 0.366140, 0.376250, 0.385310, 0.395760, 0.411070, 0.430620, 0.450040, 0.469780, 0.488820, 0.505410, 0.520690, 0.537010, 0.554650, 0.571780, 0.586870, 0.602200, 0.620460, 0.640160, 0.656310, 0.673330, 0.697510, 0.725370, 0.753600, 0.783220, 0.819860, 0.899440, 1.000000]. Each window was minimized and then equilibrated for 440 ps in the NPT ensemble, followed by 2 ns of equilibration with a heating cycle from 0 K to 300 K and then an NPT equilibration of about 8 ns. The production stage was then performed for 5 ns under NPT conditions. In this stage, hydrogen mass repartitioning was used, allowing a 4 fs timestep. A harmonic restraint centered at 2.10 Å between $Mg^{2+}$) and the phosphate oxygen was applied in a $\lambda$-dependent manner for the RNA and reference systems. This restraint was inactive at $\lambda = 0$ and smoothly activated as the particle is dummy ($k = 30$ kcal $\cdot$ mol$^{-1}$ $\cdot$ Å$^{-2}$). Window free energies were combined by MBAR[8,9] to obtain $\Delta G_{\mathrm{RNA}}$ and $\Delta G_{\mathrm{ref}}$.

### Analysis of alchemical free energy simulations

**A**

**RNA**

| Decomposition of <ΔG> into trials | | | | |
|---|---|---|---|---|
| Sampling | <ΔG> | ΔG#1 | ΔG#2 | ΔG#3 |
| | 453.30 ± 0.44 | 453.18 ± 0.60 | 452.61 ± 0.05 | 454.12 ± 0.06 |

**B**

**Reference**

| Decomposition of <ΔG> into trials | | | | |
|---|---|---|---|---|
| Sampling | <ΔG> | ΔG#1 | ΔG#2 | ΔG#3 |
| | 443.60 ± 0.40 | 443.34 ± 0.50 | 444.32 ± 0.04 | 443.01 ± 0.05 |

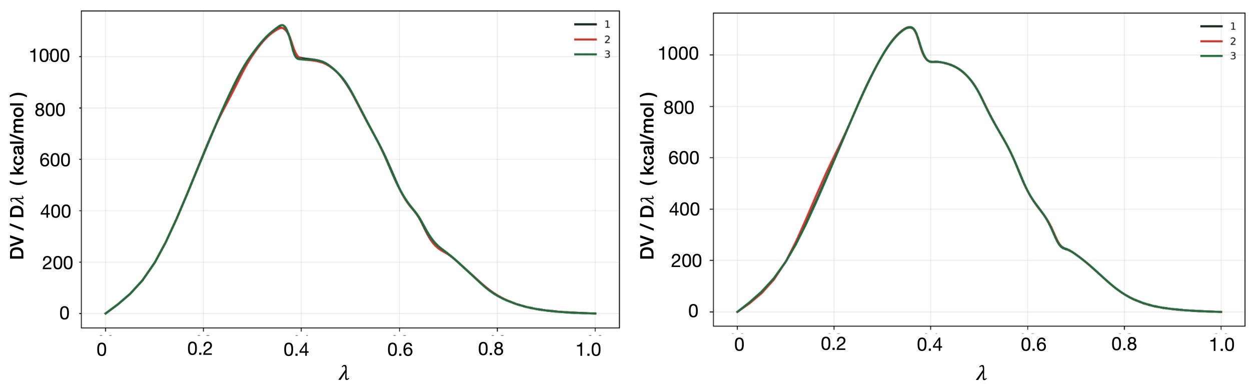


Figure S2: Analysis of alchemical free energy simulations for ion decoupling from two distinct environments: the RNA-SAM complex (A) and the phosphate reference system (B). Average $\frac{dU}{d\lambda}$ profiles obtained by MBAR[8] over three independent trials are shown. The cubic spline fit was integrated to determine the free energy of ion decoupling in each system.

### Quantum Mechanics (QM)/Molecular Mechanics (MM) Simulations

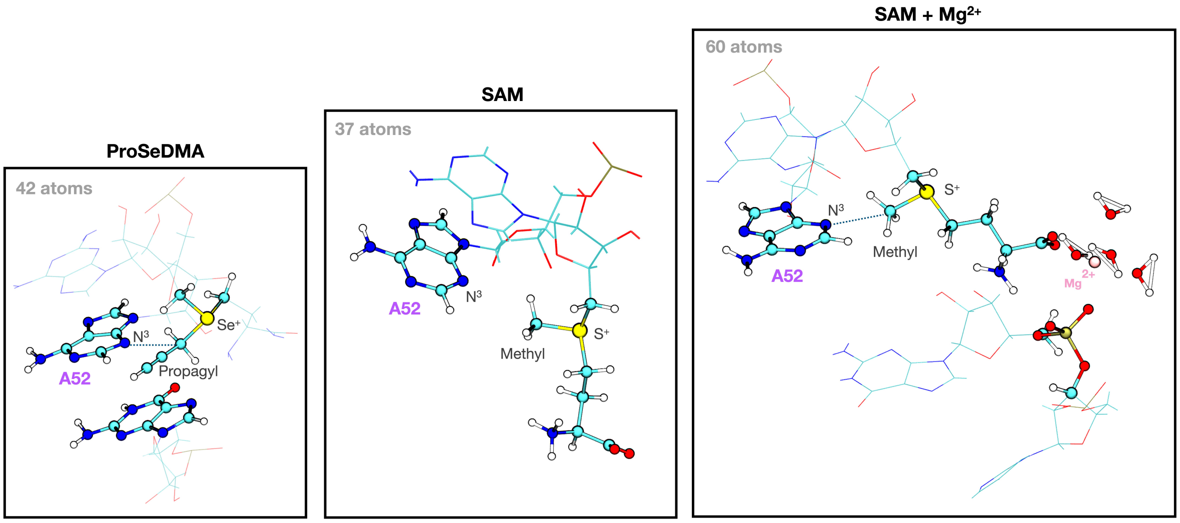


Figure S3: Representation of the different QM regions used (atoms shown with CPK representation) for the ProSeDMA (left), SAM (center), and SAM-$Mg^{2+}$ (right) reactions.

#### *ab initio* p$K_a$ calculation

The c[1]A:52 and c[7]A:52 modifications were investigated by QM/MM simulations based on the hypothesis that replacing an endocyclic nitrogen, an electron-withdrawing atom, with carbon would favor the localization of negative charge at the N3 position. This was expected to enhance nucleophilicity and, consequently, increase the p$K_a$. A recent study reported predicted p$K_a$ values for related modified nucleobases,[10] but it underestimates the N3 p$K_a$ of adenine (0.8 vs. 1.5 experimentally[11]). We therefore recomputed these p$K_a$ values and developed a more reliable predictive model leveraging available experimental data. To this end, a series of nitrogen-containing bases (Table S3) with experimentally measured p$K_a$ values was selected, and deprotonation free energies were calculated as described in the Computational Methods. These data were then used to fit a regression model for predicting experimentally unknown p$K_a$ values.

In this study, the neutral form of adenine (A) is defined as the N1, N3, and N7 deprotonated state, while the A:N1 is defined as the protonated form that carries a positive charge on the N1 site. To ensure consistency, reference compounds with the same protonation pattern—namely, neutral deprotonated structures and positively charged protonated states with known experimental p$K_a$ values were selected as benchmarks for p$K_a$ prediction, with results given in Table S3.

A linear regression analysis was performed on these nine data points, yielding the relationship: $y = 0.671x + 2.021$, with $R^2 = 0.9626$, where y represents the experimental p$K_a$ and x represents the calculated p$K_a$. The resulting fitting curve is shown in Figure S4, demonstrating a strong linear correlation between computed and experimental p$K_a$ values.

To further evaluate the predictive accuracy of this model, six additional compounds with known experimental p$K_a$ values exhibiting analogous protonation patterns—specifically, neutral deprotonated and positively charged protonated states were examined, including Purine, 2-aminopurine (2AP), 2,6-diaminopurine (DAP), cytidine (C), 5-methylcytidine (5mC), and 5-azacytidine (5nC). The same QM computational protocol was applied, and p$K_a$ values were

**Table S3: Predicted p$K_a$'s of benchmark modified nucleobases from *ab initio* DFT calculations**

| Compound | Position | $pK_a^{calc}$ | $pK_a^{expt}$ | $pK_a^{pred}$ | $pK_a{}^{Error}$ |
|---|---|---|---|---|---|
| A | N1 | 1.84 | 3.63[11] | 3.25 | -0.38 |
| A | N3 | 0.28 | 1.50[11] | 2.21 | 0.71 |
| A | N7 | -0.30 | 2.15[11] | 1.82 | -0.33 |
| $c^3$A | N1 | 6.74 | 6.80[12] | 6.55 | -0.25 |
| $c^7$A | N1 | 4.17 | 5.30[12] | 4.82 | -0.48 |
| $n^8$A | N1 | 0.27 | 2.20[13] | 2.20 | 0.00 |
| $m^6$A | N1 | 3.29 | 4.01[14] | 4.23 | 0.22 |
| $m^{66}$A | N1 | 3.96 | 4.50[14] | 4.68 | 0.18 |
| $m^7$G | N1 | 8.21 | 7.2[15] | 7.53 | 0.33 |

p$K_a$ values are calculated (calc), taken from the experimental literature (expt), and predicted (pred) as indicated by superscripts. The error indicates the difference between the experimental and predicted p$K_a$s. Including those of 8-azaadenosine ($n^8$A), 6-methyladenosine ($m^6$A), N6,N6-dimethyladenosine ($m^{66}$A), and N7-methylguanosine ($m^7$G).

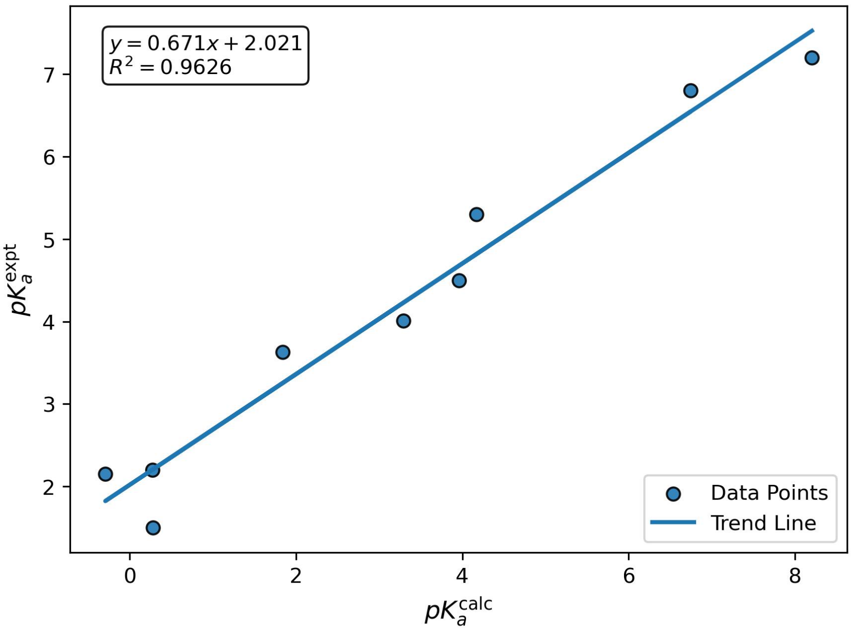


Figure S4: Experimental p$K_a$ (p$K_a{}^{expt}$) vs. calculated p$K_a$ (p$K_a{}^{calc}$)

predicted using the established regression equation. The prediction error ($\Delta$p$K_a{}^{pred-exp}$), defined as the difference between predicted and experimental p$K_a$ values, was found to be less than 1 p$K_a$ unit for all test cases, indicating good reliability and transferability of the model.

**Table S4: Calculated and experimental p$K_a$ values and predicted results.**

| Compound | Position | $pK_a^{\text{calc}}$ | $pK_a^{\text{expt}}$ | $pK_a^{\text{pred}}$ | $\text{p}K_a{}^{\text{Error}}$ |
|---|---|---|---|---|---|
| Purine | N1 | 0.81 | 2.40[16] | -1.03 | 0.16 |
| 2AP | N1 | 2.73 | 3.80[17] | 0.89 | 0.05 |
| DAP | N1 | 4.22 | 5.10[17] | 2.38 | -0.25 |
| C | N3 | 2.61 | 4.20[18] | 0.77 | -0.43 |
| 5nC | N3 | -0.48 | 2.64[19] | -2.32 | -0.94 |
| 5mC | N3 | 3.37 | 4.30[20] | 1.53 | -0.02 |

p$K_a$ values are calculated (calc), taken from the experimental literature (expt), and predicted (pred) as indicated by superscripts. The error indicates the difference between the experimental and predicted p$K_a$s. Compounds include Purine, 2-aminopurine (2AP), 2,6-diaminopurine (DAP), cytidine (C), 5-methylcytidine (5mC), and 5-azacytidine (5nC).

Here, the known experimental reference value of 1.5[11] (i.e., $\text{p}K_a^{\text{expt, A:N3}}$) is used to obtain the difference with the predicted p$K_a$ value ($\text{p}K_a^{\text{pred, A:N3}}$) and to derive $\text{p}K_a^{\text{Error}}$ (Equation 3) to correct the predicted p$K_a$ values for $c^1$A and $c^7$A at the N3 position (Equation 4).

With the validated function from our model, the predicted p$K_a$ values at the N3 site are 5.20, 4.03, and 2.21 for $c^1$A, $c^7$A, and adenine (A:N3), respectively (Table S4). The $\text{p}K_a^{\text{Error}}$ for A:N3, for which an experimental reference is available, is 0.71. Since the p$K_a$ shift at the same site across different structures is expected to be minimal from a chemical standpoint, these values were further corrected using the same deviation, hereafter referred to as p$K_a$, yielding final values of 4.49 and 3.32 for $c^1$A:N3 and $c^7$A:N3, respectively.

$$\text{p}K_a^{\text{Error}} = \text{p}K_a^{\text{pred,A:N3}} - \text{p}K_a^{\text{expt, A:N3}} \tag{3}$$

$$\text{p}K_a = \text{p}K_a^{\text{pred}} - \text{p}K_a^{\text{Error}} \tag{4}$$

### Structure and dynamics of SAMURI in solution

#### Global stability of the solvated ribozyme

##### Root Mean Square Deviation (RMSD)

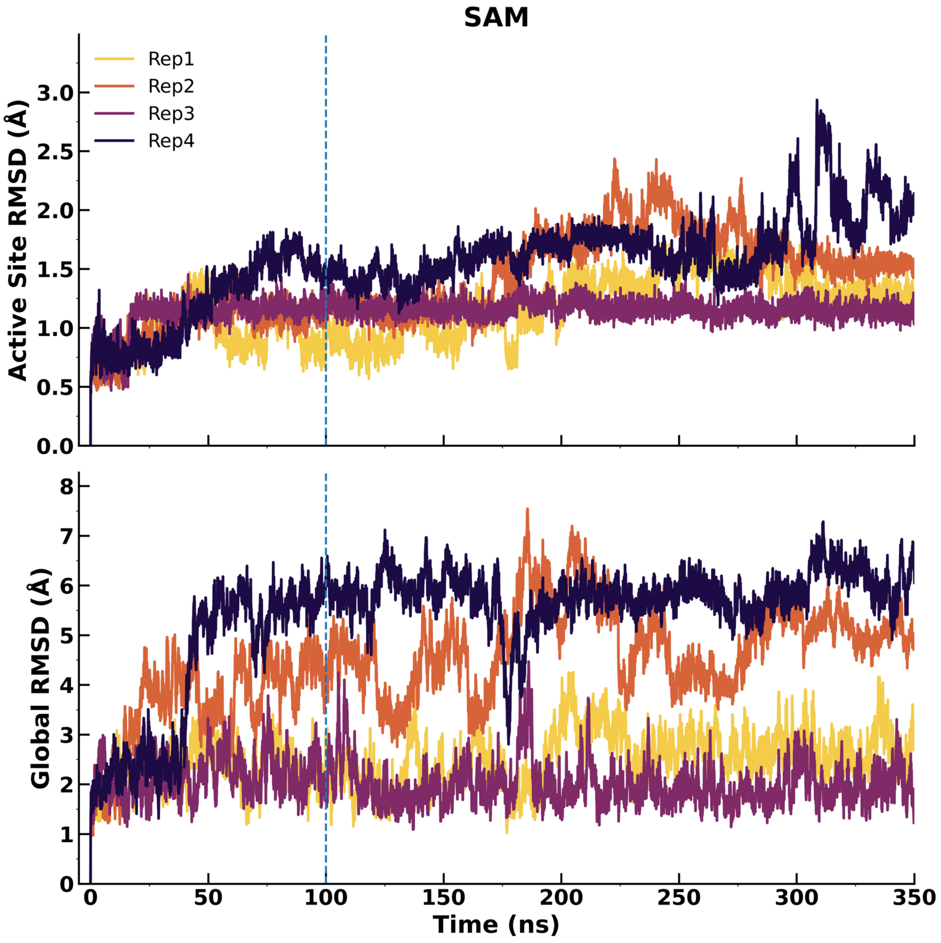


Figure S5: RMSD time series for the four replicas of SAM for the active-site residues (residues 9–15, 30–37, 52 and 62) and the global system, across the four independent replicas.

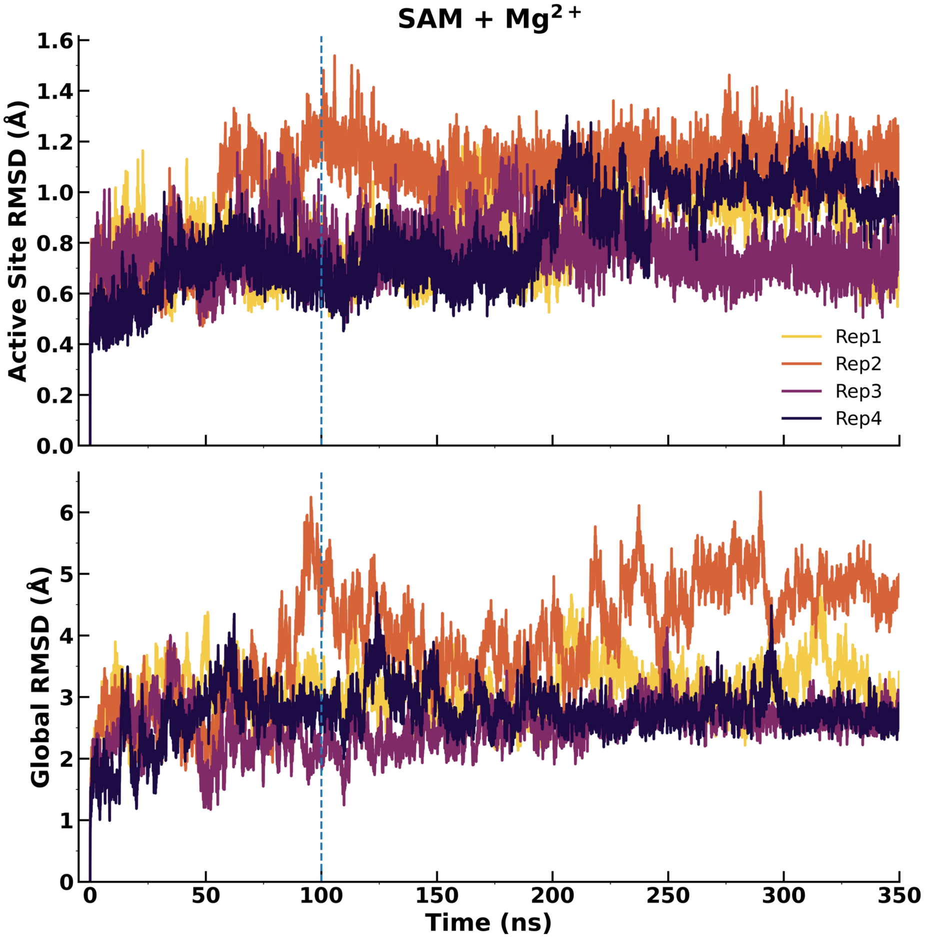


Figure S6: RMSD time series for the four replicas of SAM interacting with $Mg^{2+}$ for the active-site residues (residues 9–15, 30–37, 52 and 62) and the global system, across the four independent replicas.

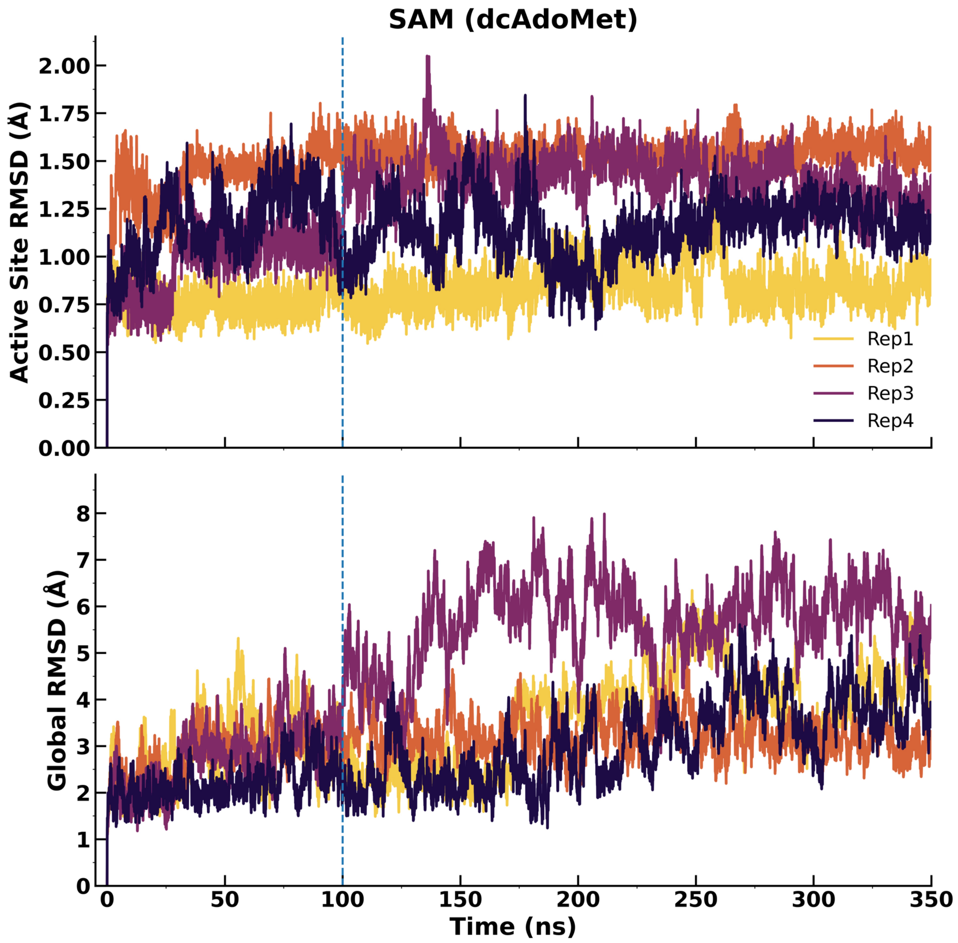


Figure S7: RMSD time series for the four replicas of SAM dcAdoMet for the active-site residues (residues 9–15, 30–37, 52 and 62) and the global system, across the four independent replicas.

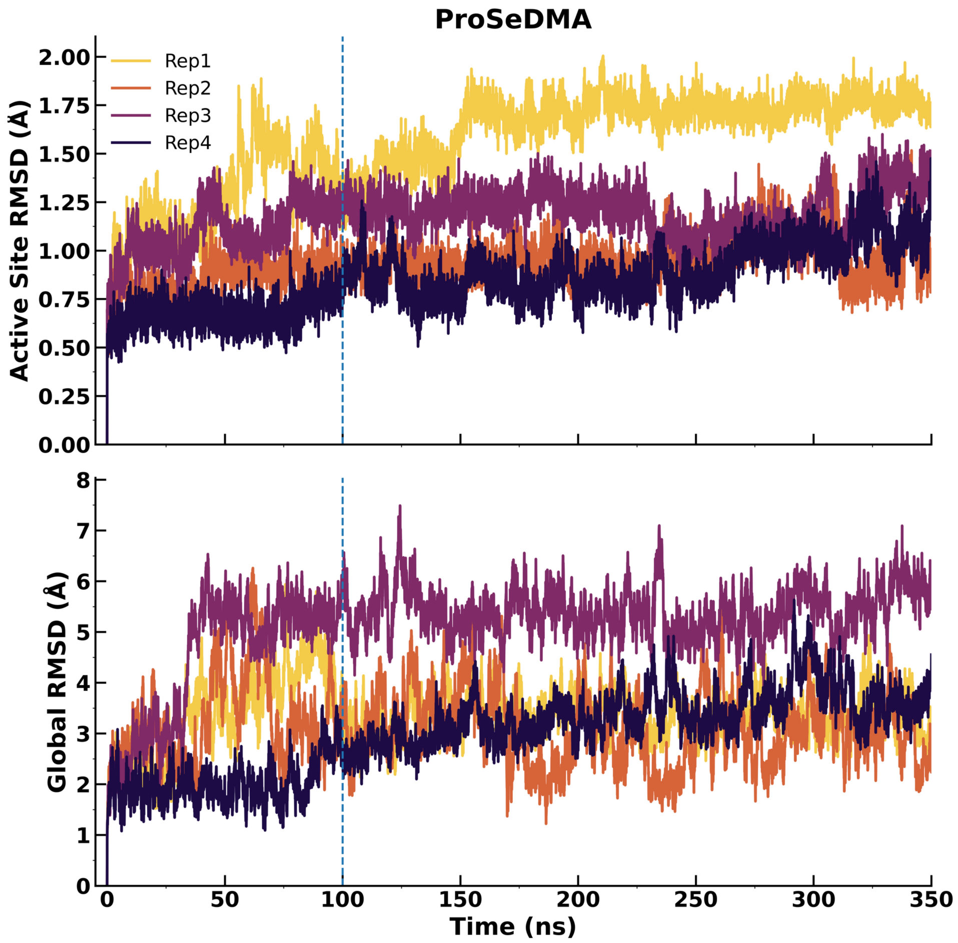


Figure S8: RMSD time series for the four replicas of ProSeDMA for the active-site residues (residues 9–15, 30–37, 52 and 59) and the global system, across the four independent replicas

#### Root Mean Square Fluctuation (RMSF)

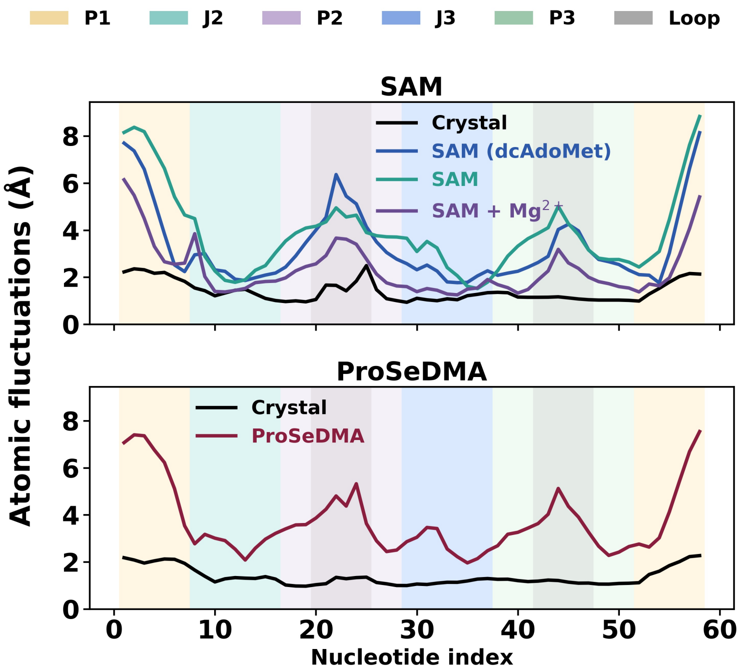


Figure S9: Root mean square atomic fluctuations (RMSF, Å) as a function of nucleotide index for the SAMURI ribozyme. Top: Systems containing SAM. The black curve represents the crystal structure. Fluctuations are shown for SAM dcAdoMet (blue), SAM (green), and SAM in the presence of $Mg^{2+}$ (purple). Bottom: System containing ProSeDMA. The black curve represents the RMSF obtained from crystal structure B-factors and the burgundy curve corresponds to the ProSeDMA-bound form. Shaded regions indicate structural domains: P1, J2, P2, J3, P3 and the loop.

### Time series and joint distributions for all simulated systems

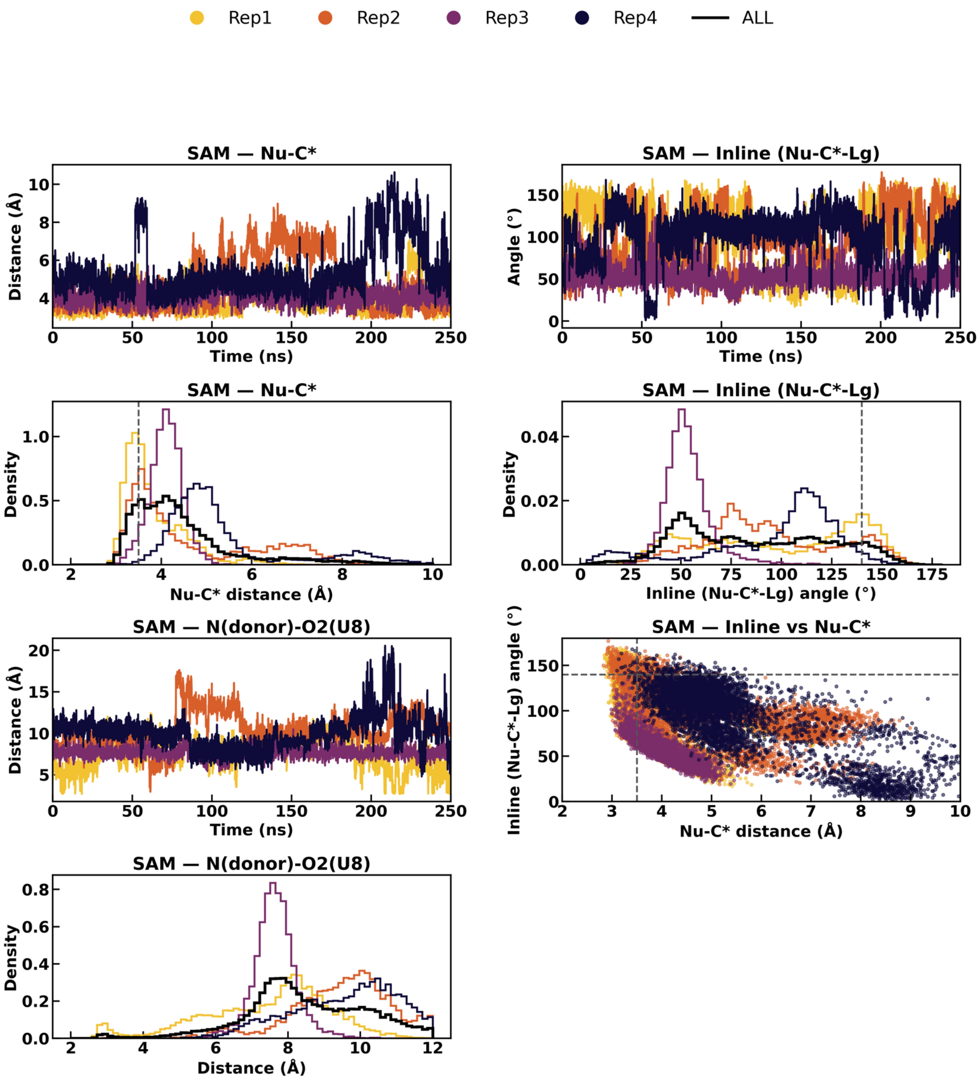


Figure S10: Time evolution of the Nu-C* distance, the inline attack angle (Nu-C*-Lg), and the N(donor)-O2(U8) distance, together with the joint distribution of the Nu-C* distance and inline angle, for the four independent 250 ns replicas of SAM.

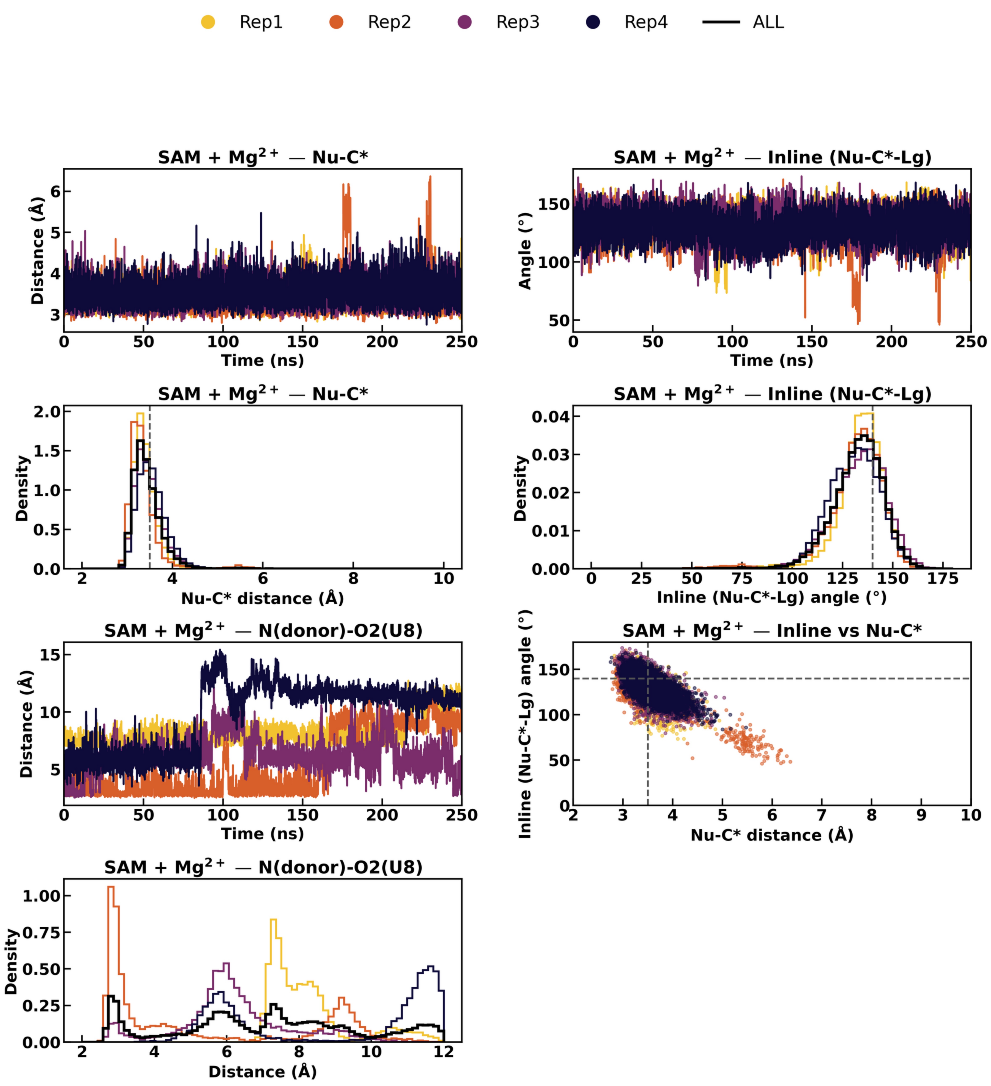


Figure S11: Time evolution of the Nu-C* distance, the inline attack angle (Nu-C*-Lg), and the N(donor)-O2(U8) distance, together with the joint distribution of the Nu-C* distance and inline angle, for the four independent 250 ns replicas of SAM in the presence of $Mg^{2+}$.

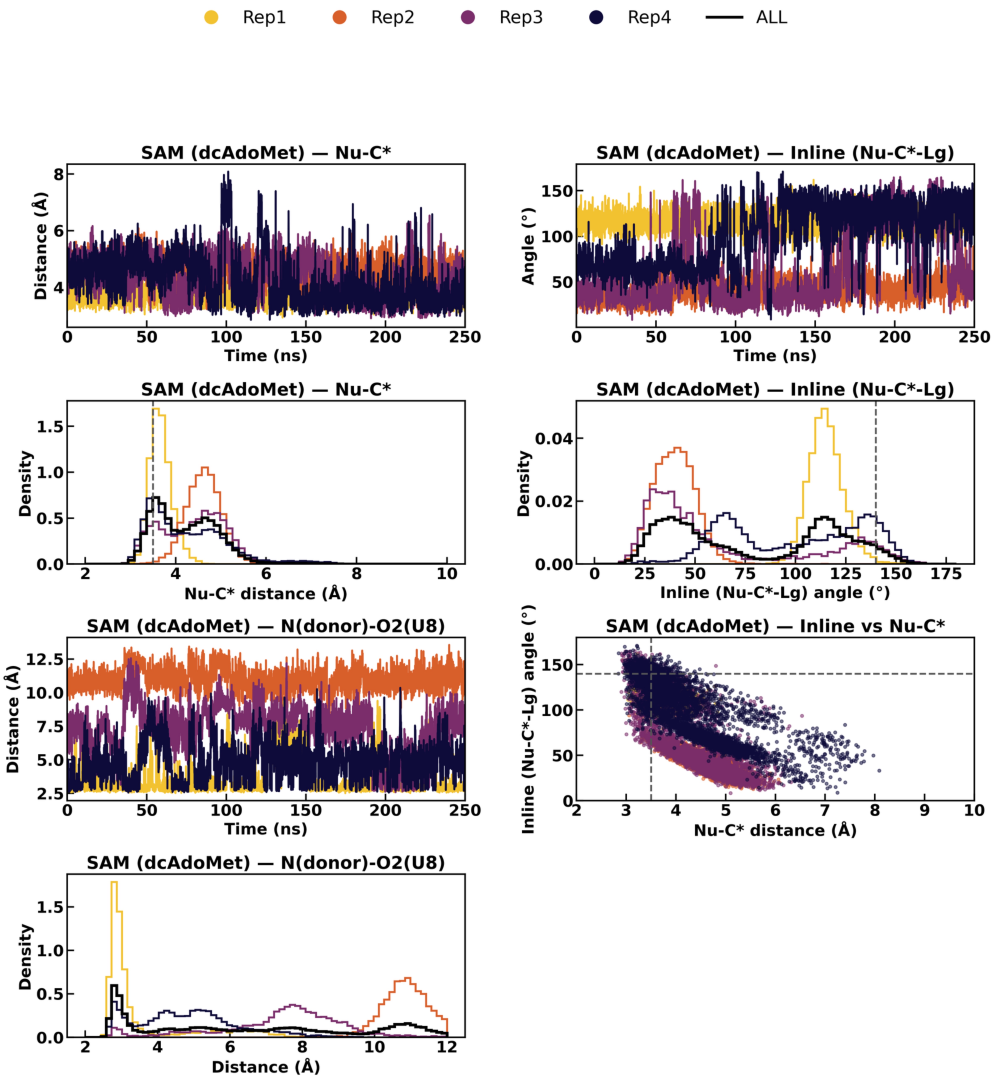


Figure S12: Time evolution of the Nu-C* distance, the inline attack angle (Nu-C*-Lg), and the N(donor)-O2(U8) distance, together with the joint distribution of the Nu-C* distance and inline angle, for the four independent 250 ns replicas of SAM (dcAdoMet).

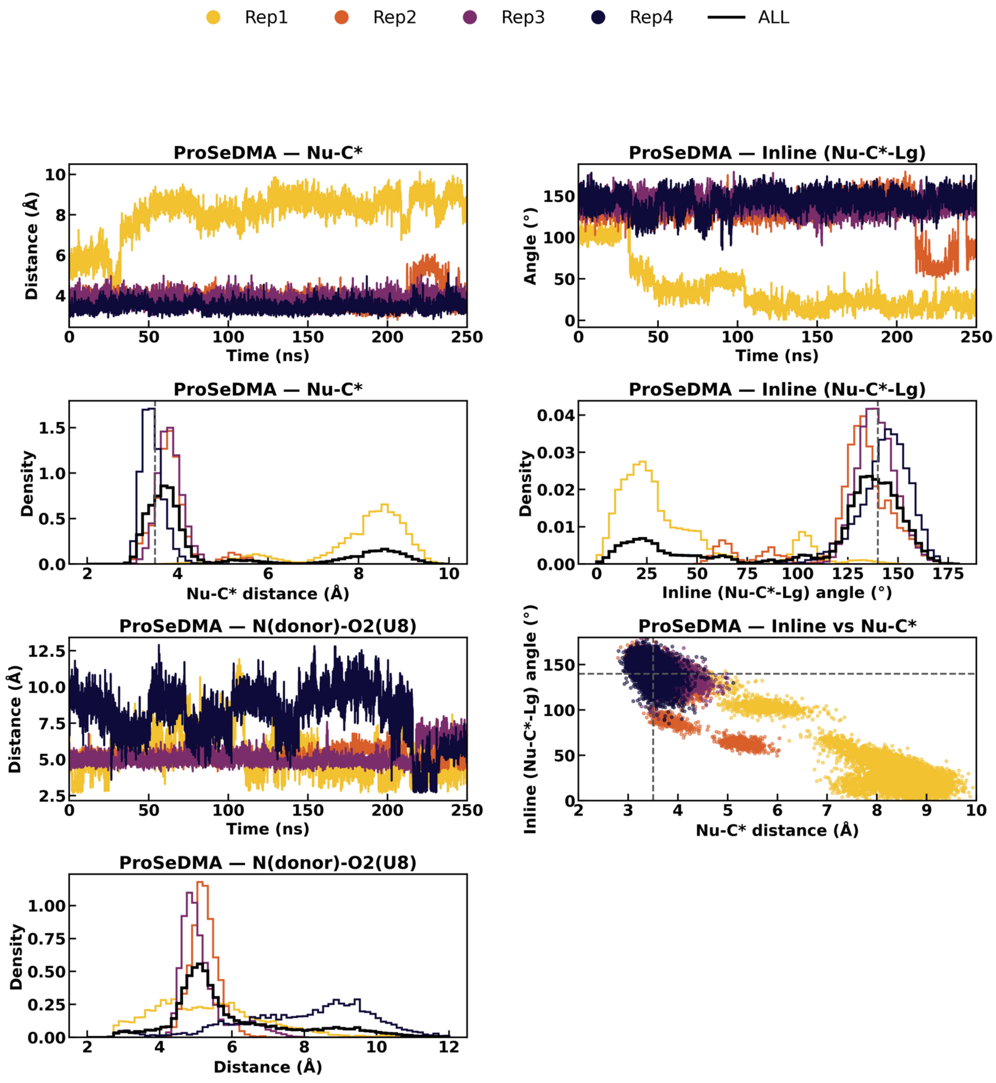


Figure S13: Time evolution of the Nu-C* distance, the inline attack angle (Nu-C*-Lg), and the N(donor)-O2(U8) distance, together with the joint distribution of the Nu-C* distance and inline angle, for the four independent 250 ns replicas of ProSeDMA.

#### Magnesium coordination in cofactor binding

In addition to the $Mg^{2+}$ ion coordinated to the cofactor tail described in the main text (shown in orange in Figure S15A), we identified the need for a further divalent ion bridging the pro-$R_P$ oxygens of residues U12 and C11 (highlighted in green Figure S15A). This site was first suggested by the accumulation of $Na^+$ ions at short distances in this region (Figure S15B) and was further supported by a pronounced $Mg^{2+}$ density maximum in the 3D-RISM analysis (Figure S15C). The completed SAM model therefore contains seven $Mg^{2+}$ ions in total.

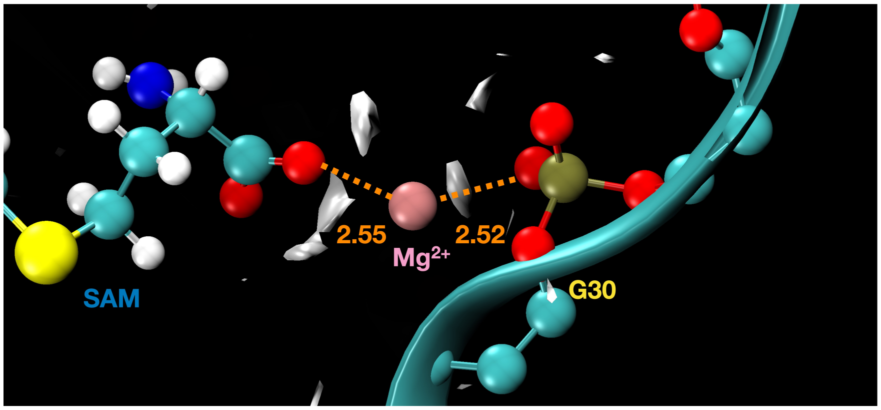


Figure S14: Prediction of a $Mg^{2+}$ binding site from 3D-RISM from the crystal structure. A density maximum was identified between the SAM carboxylate and the G30 phosphate. A $Mg^{2+}$ ion was therefore placed at ∼2.5 Å from the pro-$S_P$ oxygen of G30 and from a SAM carboxylate oxygen. This chemically plausible starting geometry was subsequently refined during equilibration.

However, the structure with the ProSeDMA cofactor does not have a carboxylate group at the end of the cofactor and therefore does not have the additional $Mg^{2+}$ observed in the SAM system. Nevertheless, 3D-RISM analysis revealed a comparable $Mg^{2+}$ density between the pro-$R_P$ oxygens of U12 and C11, and here too, a divalent ion was placed in direct coordination with these phosphates (shown in green in Figure S15A). The corresponding Mg–O distances are slightly longer (∼2.7 Å). The ProSeDMA model therefore contains four $Mg^{2+}$ ions in total. Figures S15B and S15C highlight this location by comparing the $Na^+$ artifacts observed in the absence of $Mg^{2+}$ with the corresponding $Mg^{2+}$ density predicted by 3D-RISM.

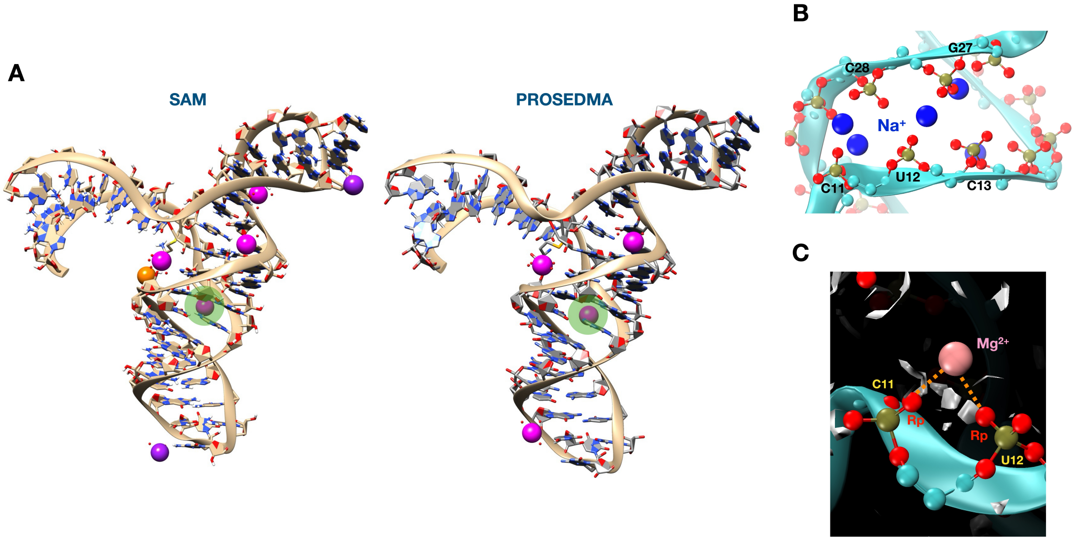


Figure S15: **(A)** Representations of SAMURI ribozyme with SAM (left) and ProSeDMA (right) cofactors, with $Mg^{2+}$ ions shown as spheres. For SAM, two $Mg^{2+}$ ions shown in purple correspond to those most likely associated with crystal packing. Because our simulations contain only a single subunit, these ions had to be restrained in order to be maintained at their crystallographic positions. The $Mg^{2+}$ bound near the cofactor carboxylate in orange, and the additional bridging ion (U12–C11) in green. For ProSeDMA, the analogous bridging $Mg^{2+}$ is shown in green. **(B)** Snapshot showing accumulation of $Na^{+}$ ions of clustered negative charges in the absence of $Mg^{2+}$. **(C)** 3D-RISM prediction from the crystal of $Mg^{2+}$ density, supporting placement of a bridging ion between pro-$R_P$ oxygens of U12 and C11. Distances between $Mg^{2+}$ and the pro-$R_P$ oxygens are ∼2.5 Å in SAM and ∼2.7 Å in ProSeDMA.

#### ProSeDMA derivatives

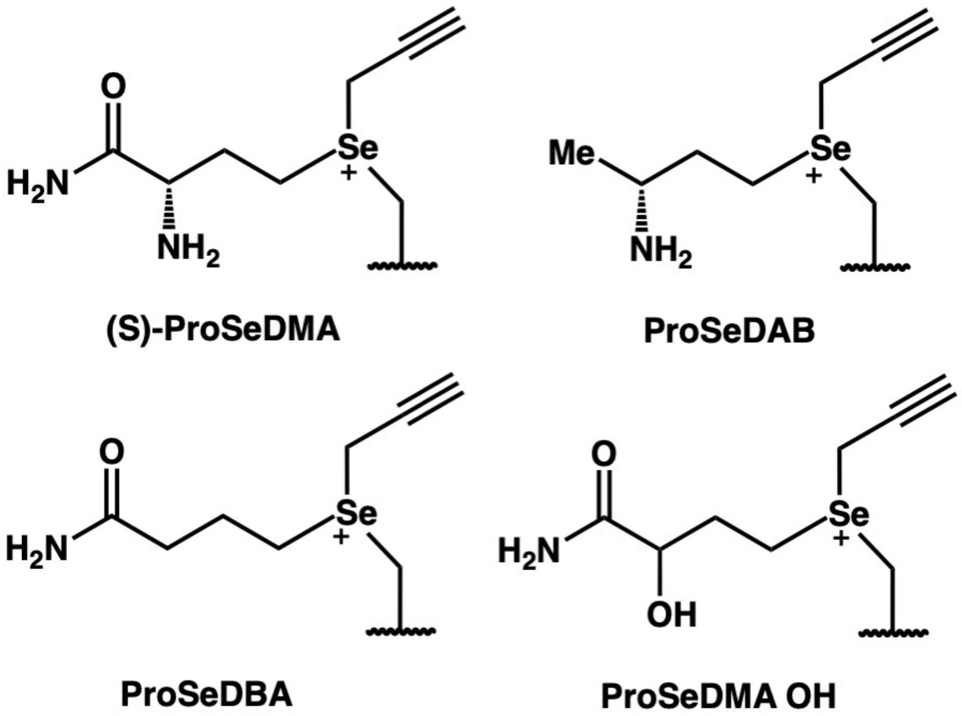


Figure S16: Chemical structure of ProSeDMA methionine unit and its derivatives mentioned in the main text. S-ProSeDMA: S-propargylic Se-2,6-diaminopurin-ribosyl-selenomethionineamide, ProSeDAB: Propargylic Se-2,6-Diaminopurineribosylseleno-2-(R)-amino-butane,ProSeDBA: Se-Propargyl-Se-2,6-Diaminopurineribosyl-selenobutanamide and ProSeDMA OH: Se-Propargyl-Se-2,6-diaminopurineribosyl-2-hydroxy-4-selenobutanamide.

#### Depurination pathway of c¹A variants

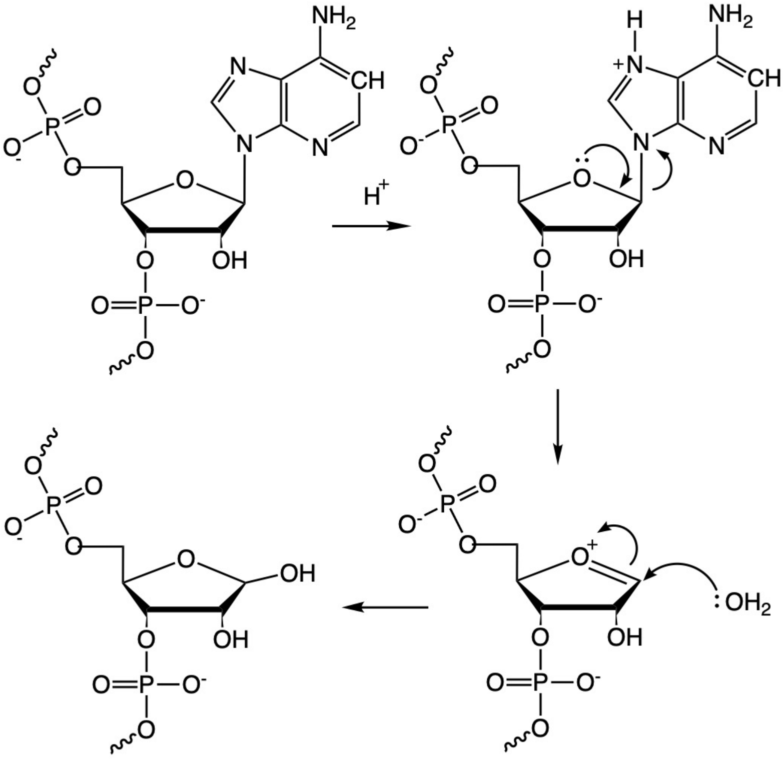


Figure S17: Depurination mechanism of adenosine resulting in the minor depurinated product for reactions of the c¹A modified ribozyme.

#### Examples of SAM-binding RNA motifs with divalent ions

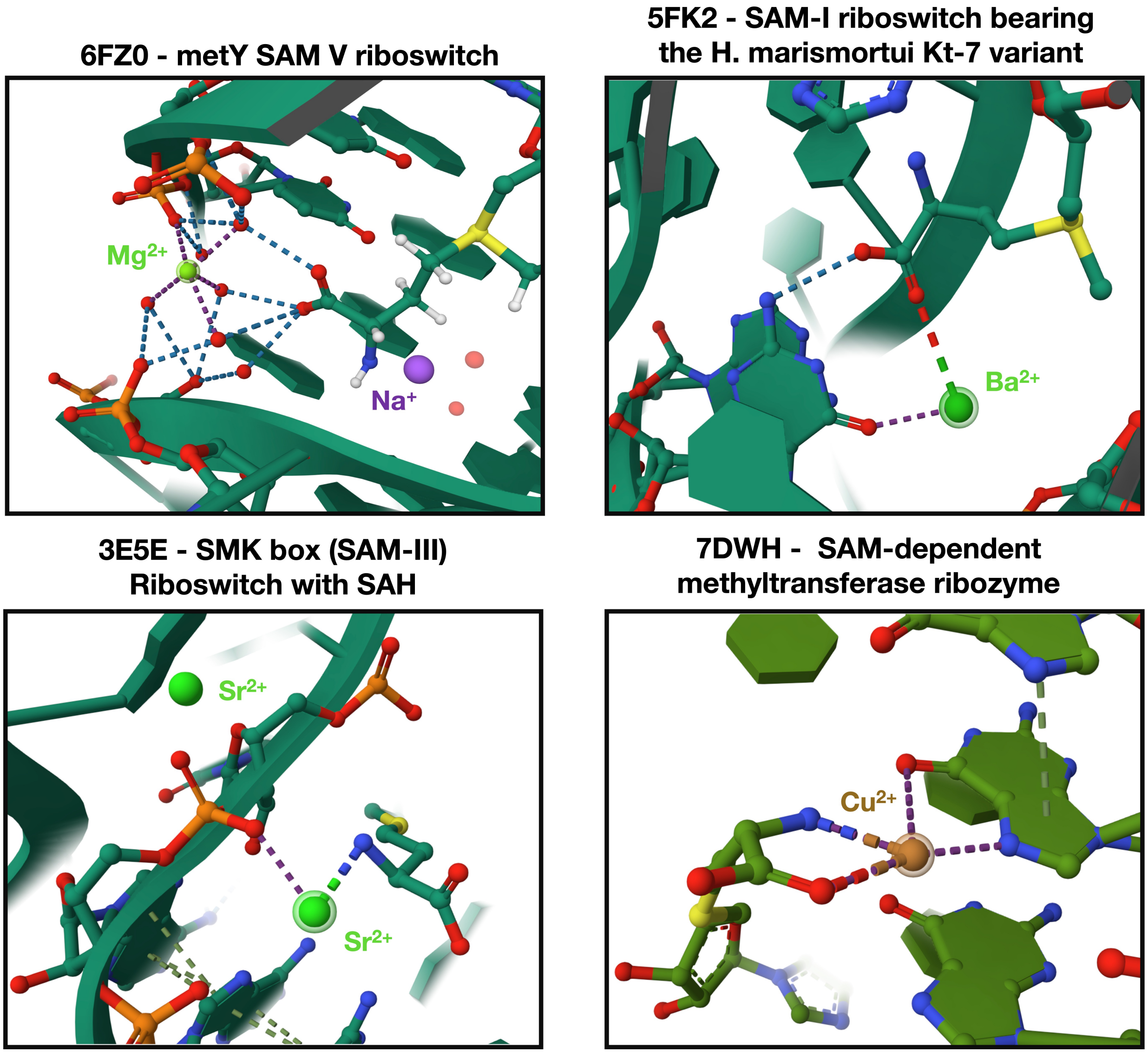


Figure S18: Representation of examples illustrating systems in which SAM acts as a cofactor interacting with divalent ions. The PDB codes are mentioned.[21–24]

**Table S5: Main interaction partners of the cofactor amine nitrogen(s) in the different systems. Fractions indicate the proportion of frames in which the corresponding contact is observed. For ProSeDMA, the two amine nitrogens are reported separately; N05 corresponds to the N$\alpha$.**

**SAM**

| Atom | Partner | Fraction |
|---|---|---|
| :62@N1 | G53@O5′ | 0.24 |
| :62@N1 | G53@OP2 | 0.13 |
| :62@N1 | G9@OP1 | 0.13 |
| :62@N1 | A7@N7 | 0.12 |
| :62@N1 | G10@OP2 | 0.09 |
| :62@N1 | G9@OP2 | 0.07 |
| :62@N1 | G53@O4′ | 0.04 |
| :62@N1 | G53@O6 | 0.04 |
| :62@N1 | A52@O3′ | 0.03 |
| :62@N1 | U8@OP2 | 0.02 |
| :62@N1 | G10@OP1 | 0.01 |
| :62@N1 | U5@O4 | 0.01 |

**SAM +Mg$^{2+}$**

| Atom | Partner | Fraction |
|---|---|---|
| :62@N1 | G10@OP1 | 0.69 |
| :62@N1 | A7@N7 | 0.31 |
| :62@N1 | A7@N1 | 0.17 |
| :62@N1 | G10@OP2 | 0.12 |
| :62@N1 | A7@N6 | 0.07 |
| :62@N1 | U8@O2 | 0.05 |
| :62@N1 | U8@O4 | 0.05 |
| :62@N1 | G53@O6 | 0.03 |

**SAM (dcAdoMet)**

| Atom | Partner | Fraction |
|---|---|---|
| :62@N1 | G10@OP1 | 0.22 |
| :62@N1 | G53@O5′ | 0.08 |
| :62@N1 | G53@O4′ | 0.08 |
| :62@N1 | G10@OP2 | 0.05 |
| :62@N1 | G53@OP2 | 0.05 |
| :62@N1 | U8@O2 | 0.04 |
| :62@N1 | A52@O3′ | 0.03 |
| :62@N1 | G53@O6 | 0.03 |
| :62@N1 | A7@N7 | 0.03 |
| :62@N1 | G10@O6 | 0.02 |
| :62@N1 | U8@O2′ | 0.01 |
| :62@N1 | G53@N7 | 0.01 |

**ProSeDMA**

| Atom | Partner | Fraction |
|---|---|---|
| *N05 (N$\alpha$)* | | |
| :52@N05 | G53@O4′ | 0.28 |
| :52@N05 | G53@O5′ | 0.21 |
| :52@N05 | G10@N7 | 0.16 |
| :52@N05 | G9@O2′ | 0.10 |
| :52@N05 | U8@O2′ | 0.07 |
| :52@N05 | A52@O3′ | 0.07 |
| :52@N05 | G53@OP2 | 0.05 |
| :52@N05 | U8@O2 | 0.02 |
| :52@N05 | A52@N3 | 0.01 |
| *N01* | | |
| :52@N01 | A7@N3 | 0.24 |
| :52@N01 | G53@O5′ | 0.06 |
| :52@N01 | G53@OP2 | 0.03 |
| :52@N01 | G10@O4′ | 0.03 |
| :52@N01 | G9@O2′ | 0.03 |
| :52@N01 | A7@N7 | 0.03 |
| :52@N01 | G10@N7 | 0.01 |
| :52@N01 | G10@N9 | 0.01 |
| :52@N01 | U8@O2 | 0.01 |

### References


(1) Case, D. A.; Cerutti, D. S.; Cruzeiro, V. W. D.; Darden, T. A.; Duke, R. E.; Ghazimirsaeed, M.; Giambaşu, G. M.; Giese, T. J.; Götz, A. W.; Harris, J. A. et al. Recent Developments in Amber Biomolecular Simulations. *J. Chem. Inf. Model.* **2025**, *65*, 7835–7843.

(2) Wang, J.; Cieplak, P.; Kollman, P. A. How well does a restrained electrostatic potential (RESP) model perform in calculating conformational energies of organic biological molecules. *J. Comput. Chem.* **2000**, *21*, 1049–1074.

(3) Case, D. A.; Betz, R. M.; Cerutti, D. S.; Cheatham III, T. E.; Darden, T. A.; Duke, R. E.; Giese, T. J.; Gohlke, H.; Goetz, A. W.; Homeyer, N. et al. AMBER 16. University of California, San Francisco: San Francisco, CA, 2016.

(4) Møller, C.; Plesset, M. S. Note on an approximation treatment for many-electron systems. *Phys. Rev.* **1934**, *46*, 618–622.

(5) Loncharich, R. J.; Brooks, B. R.; Pastor, R. W. Langevin dynamics of peptides: the frictional dependence of isomerization rates of N-acetylalanyl-N'-methylamide. *Biopolymers* **1992**, *32*, 523–535.

(6) Åqvist, J.; Wennerström, P.; Nervall, M.; Bjelic, S.; Brandsdal, B. O. Molecular dynamics simulations of water and biomolecules with a Monte Carlo constant pressure algorithm. *Chem. Phys. Lett.* **2004**, *384*, 288–294.

(7) Lee, T.-S.; Lin, Z.; Allen, B. K.; Lin, C.; Radak, B. K.; Tao, Y.; Tsai, H.-C.; Sherman, W.; York, D. M. Improved Alchemical Free Energy Calculations with Optimized Smoothstep Softcore Potentials. *J. Chem. Theory Comput.* **2020**, *16*, 5512–5525.

(8) Shirts, M. R.; Chodera, J. D. Statistically optimal analysis of samples from multiple equilibrium states. *J. Chem. Phys.* **2008**, *129*, 124105.

(9) Giese, T. J.; York, D. M. FE-ToolKit: The free energy analysis toolkit. https://gitlab.com/RutgersLBSR/fe-toolkit.

(10) Mlotkowski, A. J.; Schlegel, H. B.; Chow, C. S. Calculated p$K_a$ Values for a Series of Aza- and Deaza-Modified Nucleobases. *J. Phys. Chem. A* **2023**, *127*, 3526–3534.

(11) Kapinos, L. E.; Operschall, B. P.; Larsen, E.; Sigel, H. Understanding the acid-base properties of adenosine: the intrinsic basicities of N1, N3 and N7. *Chem. Eur. J.* **2011**, *17*, 8156–8164.

(12) Bereiter, R.; Himmelstoß, M.; Renard, E.; Mairhofer, E.; Egger, M.; Breuker, K.; Kreutz, C.; Ennifar, E.; Micura, R. Impact of 3-deazapurine nucleobases on RNA properties. *Nucleic Acids Res.* **2021**, *49*, 4281–4293.

(13) Wierzchowski, J.; Wielgus-Kutrowska, B.; Shugar, D. Fluorescence emission properties of 8-azapurines and their nucleosides, and application to the kinetics of the reverse synthetic reaction of purine nucleoside phosphorylase. *Biochim. Biophys. Acta* **1996**, *1290*, 9–17.

(14) Martin, D. M. G.; Reese, C. B. Some aspects of the chemistry of N(1)- and N(6)-dimethylallyl derivatives of adenosine and adenine. *J. Chem. Soc. C* **1968**, *0*, 1731–1738.

(15) Lawley, P. D.; Brookes, P. Further Studies on the Alkylation of Nucleic Acids and their Constituent Nucleotides. *Biochem. J.* **1963**, *89*, 127–38.

(16) Kampf, G.; Kapinos, L. E.; Griesser, R.; Lippert, B.; Sigel, H. Comparison of the acidbase properties of purine derivatives in aqueous solution. Determination of intrinsic proton affinities of various basic sites. *J. Chem. Soc. Perkin Trans. 2* **2002**, *2*, 1320–1327.

(17) Han, J.; Burke, J. M. Model for General Acid-Base Catalysis by the Hammerhead Ribozyme: pH-Activity Relationships of G8 and G12 Variants at the Putative Active Site. *Biochemistry* **2005**, *44*, 7864–7870.

(18) Shugar, D.; Fox, J. J. Spectrophotometric studies of nucleic acid derivatives and related compounds as a function of pH. *Biochim. Biophys. Acta.* **1952**, *9*, 199–218.

(19) Notari, R. E.; DeYoung, J. L. Kinetics and Mechanisms of Degradation of the Antileukemic Agent 5-Azacytidine in Aqueous Solutions. *J. Pharm. Sci.* **1975**, *64*, 1148–57.

(20) Fox, J. J.; Van Praag, D.; Wempen, I.; Doerr, I. L.; Cheong, L.; Knoll, J. E.; Eidinoff, M. L.; Bendich, A.; Brown, G. B. Thiation of Nucleosides. II. Synthesis of 5-Methyl-2'-deoxycytidine and Related Pyrimidine Nucleosides¡sup¿1¡/sup¿. *J. Am. Chem. Soc.* **1959**, *81*, 178–187.

(21) Huang, L.; Lilley, D. M. Structure and ligand binding of the SAM-V riboswitch. *Nucleic Acids Res.* **2018**, *46*, 6869–6879.

(22) Huang, L.; Wang, J.; Lilley, D. A critical base pair in k-turns determines the conformational class adopted, and correlates with biological function. *Nucleic Acids Res.* **2016**, *44*, 5390–5398.

(23) Lu, C.; Smith, A. M.; Fuchs, R. T.; Ding, F.; Rajashankar, K.; Henkin, T. M.; Ke, A. Crystal structures of the SAM-III/SMK riboswitch reveal the SAM-dependent translation inhibition mechanism. *Nat. Struct. Mol. Biol.* **2008**, *15*, 1076–1083.

(24) Jiang, H.; Gao, Y.; Zhang, L.; Chen, D.; Gan, J.; Murchie, A. I. H. The identification and characterization of a selected SAM-dependent methyltransferase ribozyme that is present in natural sequences. *Nat Catal* **2021**, *4*, 872–881.
